# Long-term exposure to microplastics and heat affects bumblebee behavior patterns, colony development and social networks

**DOI:** 10.1101/2025.07.24.666509

**Authors:** Dong Sheng, Siyuan Jing, Marcel Balle, Thomas Cherico Wanger

**Author notes:** Corresponding authors (DS); (TCW). Contributed equally (Dong Sheng, Siyuan Jing).

## Abstract

Pollinators are crucial for terrestrial ecosystems and global food security, but their populations are declining from multiple stressors including pollution and climate change. The effects of plastic pollution alone or in interaction with climate change on pollinators remain largely unexplored. Here, we investigate sublethal microplastic exposure effects on pollinators at individual, colony and network level in combination with heating to simulate climate warming. We conducted a one-generation trial on 30 bumblebee (*Bombus terrestris*) colonies in a customized beekeeping structure with a nesting and a foraging room, where colonies were maintained under optimal temperature (25 □) or heated condition (30 □), and fed with oil-seed rape pollens and 50% w/w sucrose solution containing 0, 10, or 100 mg/L 30 μm polyethylene beads. Real-time tracking with object detection (F1 = 83%) and matrix code scanning showed that microplastic exposure and heat significantly stimulated individual activity and altered labor division. Workers shifted to nursing behaviors and foraged more frequently for sucrose solution and less for pollen. Brood development was impaired by up to 48%, and colony population growth was restrained by 12–15%. Microplastic exposure and heating also significantly intensified social interactions and increased the dominance of the queen in her colony. These findings suggest that plastic pollution has complex, cross-level impacts on bumblebee colonies and their pollination potential, which may be exacerbated under climate change.

---

Pollinators are vital to terrestrial ecosystems as up to 94% of all flowering plants benefit from animal pollination^1^, and they maintain global food security as 75% of global food production depends on pollination^2,3^. The global value of pollination service is estimated to be US$195 billion to ∼US$387 billion annually^4^, and this value will likely increase in the future with the growing proportion of agricultural area occupied by pollinator-dependent crops^5^. Despite their importance, the global population of pollinators are declining due to land use changes, pesticides, and climate change, with many of these drivers being amplified in synergy^6,7^. For instance, climate change affects pollinators by causing phenological asynchrony of plant-pollinator interactions, shifting habitable spatiotemporal ranges, and even exacerbating other drivers of decline^8^. Moreover, exposure to neonicotinoid pesticides impairs bee health, and reduces overwintering success and colony reproduction^9^. When the metabolism is enhanced by a warmer climate it can induce higher uptake of neonicotinoids and amplify resulting effects^10^. These effects likely extend to emerging contaminants such as microplastics that are only slowly being associated with pollinator decline^11^ but the interactive effects with climate change are not known.

Microplastics (hereafter MP) are persistent plastic particles ranging from 1 μm to 5 mm and accumulate in large quantities in agricultural landscapes^12^. Pollinators such as honeybees (*Apis mellifera, Apis cerana*, and *Partamona helleri*) are directly exposed to sublethal MP effects, including tissue damage in the alimentary system^13^, altered gut microbiota^14^ and gene expressions^13,14^, as well as impaired memory^15^, responsiveness^15,16^ and walking behaviors^17^. MP also make honeybees more vulnerable to other threats such as pathogens^13^ and antibiotics^14^. Current studies, however, have mainly focused on MP effects at individual level, but ignored population and colony level effects of primarily eusocial insects (including honeybees and bumblebees) in which social interactions play an important role in their survival and behavior patterns. The eusociality of many pollinators brings emerging effects at the social level, and eventually determines how individual effects lead to changes in pollination services provided by pollinator colonies. For example, a social buffer in bumblebee colonies reduces the impact of pesticides^18^. MP sampled by honeybees will be transported into beehives and finally to the larvae^19^. Yet MP effects on eusocial pollinator colonies remain unexplored to date.

Here, we address a critical knowledge gap in pollination research – the interactive effects of plastic pollution from polyethylene (hereafter PE) and climate change on pollinators at individual, colony and social level. We are using computer-vision techniques to track bumblebee (*Bombus terrestris*) responses to MP and heating in a lab experiment over one-generation to test the following four hypotheses:

1. Exposure to MP alters the specific individual behaviors: a) individual activity, b) labor division, and c) the diurnal/nocturnal rhythms in a bumblebee colony. This is, because MP may induce physiological damages to mouthparts and guts^13^ that forces individuals to behavior adaptations that can compensate for ingestion and digestion deficiency.
2. MP exposure limits bumblebee colony growth and nest structure development, because MP may be transferred to larvae, impair larvae health^19^ and alter individuals’ nest-building behaviors.
3. MP exposure shifts the social interactions of bumblebee colonies, because as eusocial insects, bumblebees may transmit information of external stressors among individuals and adopt social-level buffering^18^. This would then manifest in changes of general connectivity (hereafter centrality ranking) of individuals and especially the queen in the colony.
4. Climate change – measured as temperature increase – enhances MP effects observed in hypotheses 1 – 3, because heating greatly affects bumblebee behaviors in general^20^.

We conducted the multi-generation experiment on 30 queenright bumblebee (*Bombus terrestris*) colonies in a customized beekeeping structure with a nesting and a foraging room. Colonies were maintained under optimal or increased temperature (i.e, 25° and 30 □, respectively), and fed with oil-seed rape pollens and 50% w/w sucrose solution containing 0, 10 (low MP exposure), or 100 (high MP exposure) mg/L 30 μm PE beads. To validate successful experimental plastic exposure of the bumblebees, we measured accumulated MP particles in bumblebee samples by Raman spectra (Extended Data Fig. 1; Extended Data Fig. 2a-b and Supplementary Method 1). The number of PE MP particles in bumblebee samples significantly increased with treatment exposure levels (p < 0.05, Extended Data Fig. 1c).

**Fig. 1.**
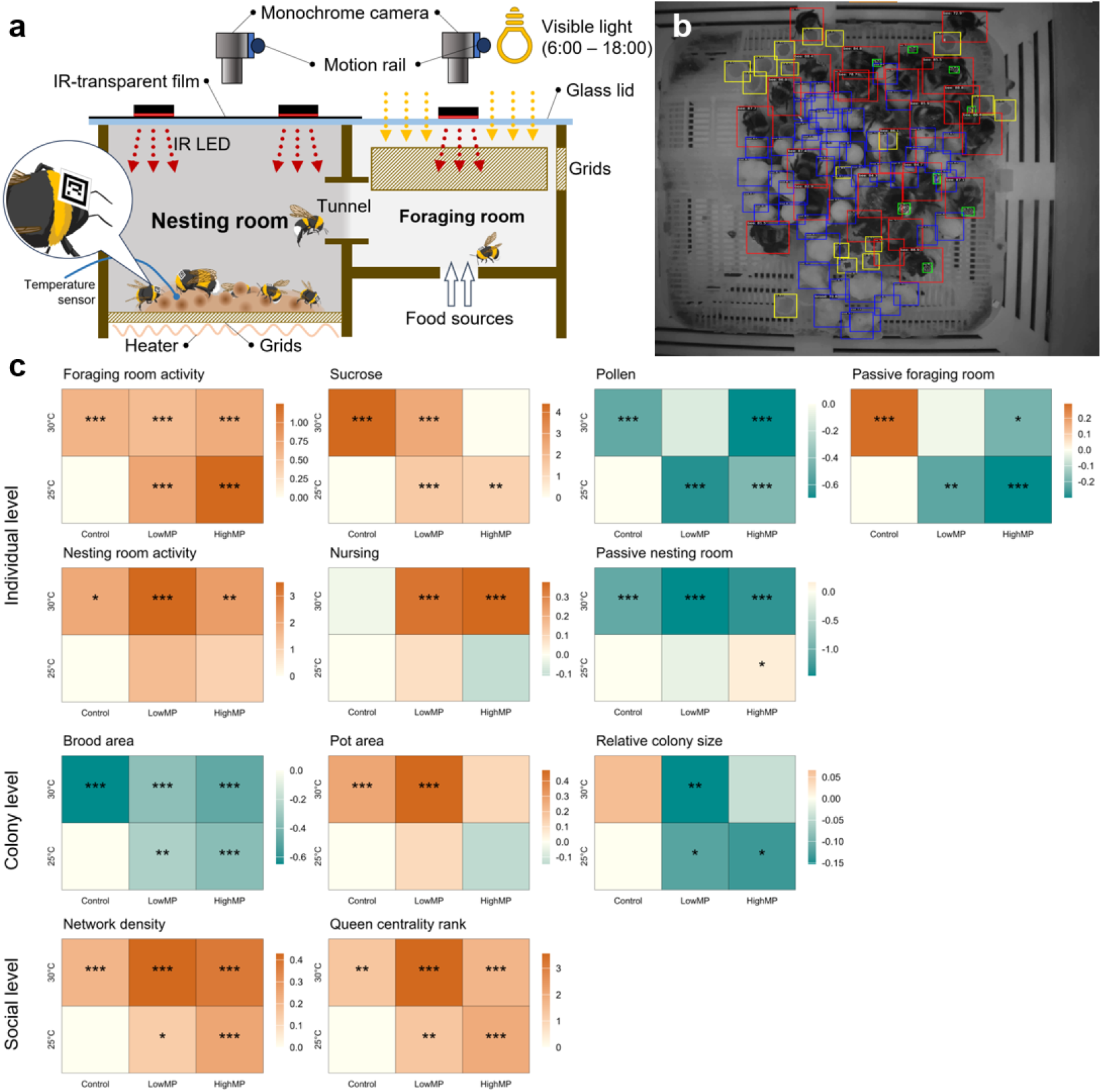
Effects of microplastic exposure and heat on bumblebees at individual, colony, and social level. **(a)** Schematic description of a bee box and monitoring system. A bee box consists of a nesting room and a foraging room, connected by a tunnel. The nesting room is kept dark within the visible light spectrum, while the foraging room follows a 12h:12h light cycle. Food sources are provided *ad libitum* through two feeders in the foraging room. Each room is illuminated by infrared LEDs and monitored by a monochrome camera. **(b)** The object detection model identifies different categories of objects from an image. Green, red, yellow and blue bounding boxes represent bee tags, bees, pots and broods respectively. **(c)** A summary of relative effect sizes for all microplastic exposure and heat treatments at colony, individual and social levels. “***”: P-value < 0.001; “**”: P-value < 0.01; “*”: P-value < 0.05. For detailed calculation see Supplementary Method 2.

**Fig. 1.**
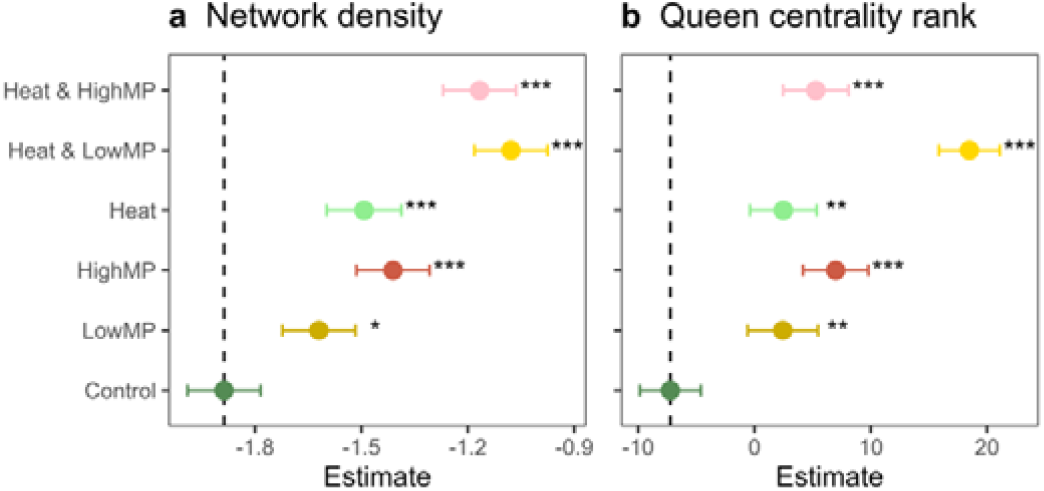
Treatment effects on the social level over time. Panel **(a)** shows the network weighted density, and **(b)** the ranking of queen centrality among all other nodes. The weighted density variable was transformed according to formula (2). “***”: P-value < 0.001; “**”: P-value < 0.01; “*”: P-value < 0.05.

## Individual behaviors

### Activity

We tagged bumblebees with miniature matrix codes^21^ and measured activity by the average moving speed of tracked individuals. Exposure to MP or elevated temperature significantly influenced bumblebee activity patterns with combined MP exposure and heat amplifying this effect. Over time, the activity of tagged individuals decreased in the foraging room and in general (Extended Data Fig. 3). As diurnal insects, bumblebees were significantly more active during the day than at night (p < 0.0001, Extended Data Fig. 4). MP exposure and/or heat significantly increased activity in the foraging room (p < 0.001) (Fig. 2a). Heat increased nesting room activity in general (p < 0.05) and especially during the day (p < 0.001) (Fig. 2e and Extended Data Fig. 7e). While MP treatments alone did not significantly alter nesting room activity, their combination with heat had a synergistic effect during the day (p < 0.001) (Fig. 2e and Extended Data Fig 7e).

**Fig. 2.**
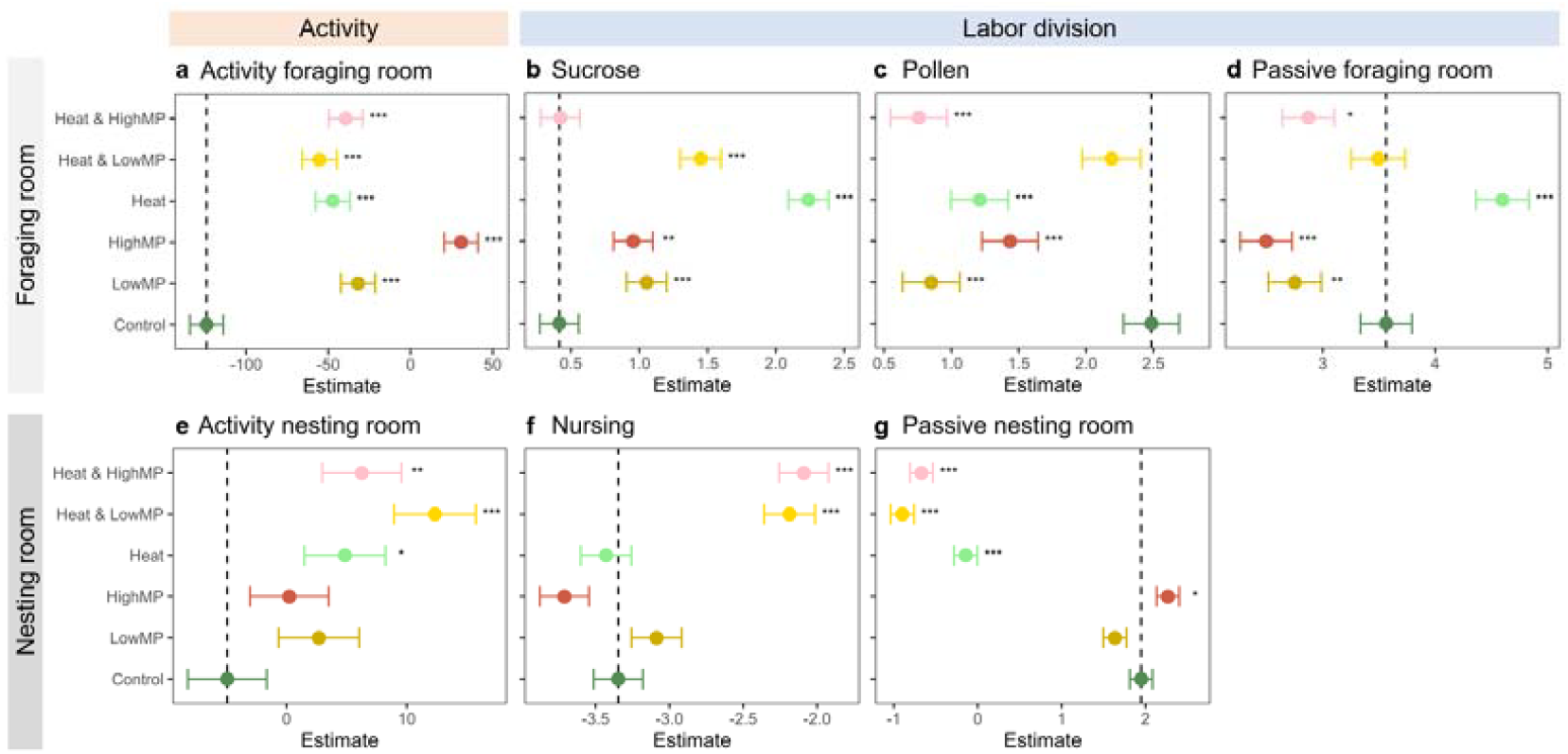
Treatment effects on the individual level over time. In the foraging room, panel **(a)** shows individual activity, while panels **(b), (c)**, and **(d)** provide details on foraging activity on sucrose solution and pollen, as well as passive behaviors in the foraging room. In the nesting room, panel **(e)** shows individual activity, while panels **(f)** and **(g)** show regression model results of nursing and passive behaviors, respectively. Labor division variables [panel **(b)** to **(d)** and **(f)** to **(g)**] were transformed according to formula. For definitions of labor division, see Extended Data Table 3. “***”: P-value < 0.001; “**”: P-value < 0.01; “*”: P-value < 0.05.

### Labor division

We classified the labor types of colony workers into five categories (Extended Data Table 3) according to the spatial locations. MP exposure and heat disrupted labor division in bumblebee colonies. Over time, we found a natural progression of more individuals shifting from nursing to foraging roles (Extended Data Fig. 5). As expected for a diurnal organism, behavioral patterns differed during day and night with the majority of individuals foraging for sucrose or pollen during the day (p < 0.0001, Extended Data Fig. 6a-b), and resting at night (p < 0.001, Extended Data Fig. 6e). For foraging related variables in combined day and night results, individual MP treatments and heat had strong significant effects, as did the combined treatment with low or high MP concentration. More visits on the sucrose feeder (p < 0.01 for high MP; p < 0.001 for low MP and/or heat) were detected, and MP effect was mainly nocturnal (Fig. 2b and Extended Data Fig. 8b). On the contrary, less visits on the pollen feeder (p < 0.001 for low MP, high MP, heat, and heat with high MP) were observed with MP exposure and heat in general and during the day (Fig. 2c and Extended Data Fig. 7c). MP exposure limited the increase of passive bees in the foraging room (p < 0.01 for low MP; p < 0.001 for high MP) especially in nocturnal times (p < 0.001), while heating urged more bees to move out especially in diurnal times (p < 0.001) (Fig. 2d and Extended Data Fig. 7d). Nursing related variables show a consistent effect of combined MP and heat effects but not individual effects in general. Specifically, nursing rate was not affected by single stressors, but combined heat and MP showed significant positive effects on nursing (Fig. 2f). The proportion of passive bees in the nesting room was positively affected by high MP (p < 0.05), but all combination treatments and heat significantly reduced passive bees (p < 0.001) suggesting that heat is the driving factor (Fig. 2e).

### Colony development

Exposure to MP and elevated temperatures significantly impacted bumblebee colony development. Brood area declined by 26% (p < 0.01) and 37% (p < 0.001) faster under low and high MP exposure, respectively (Fig. 3a). Combined exposure with heat further reduced brood area by 35% and 48% (p < 0.001), with heat alone having the strongest effect (65% reduction, p < 0.001) (Fig. 3a). The reduced additional loss in brood area with the combination of MP and heat may be the result of bumblebees putting more effort in maintaining their nest structure (Fig. 2f). These reductions suggest both independent and synergistic impacts of MP and thermal stress on colony reproduction. Moreover, pot area increased by 27% (p < 0.001) under heat and by 47% (p < 0.001) under combined heat and low MP exposure (Fig. 3b), possibly reflecting altered resource storage or stress-related compensatory behavior. Relative colony size growth declined by 12% and 13% (p < 0.05) under low and high MP exposure, respectively, and by 15% (p < 0.05) when combined with heat (Fig. 3c). No significant effects were found on the maximum colony size of bumblebee colonies (Extended Data Fig. 10).

**Fig. 3.**
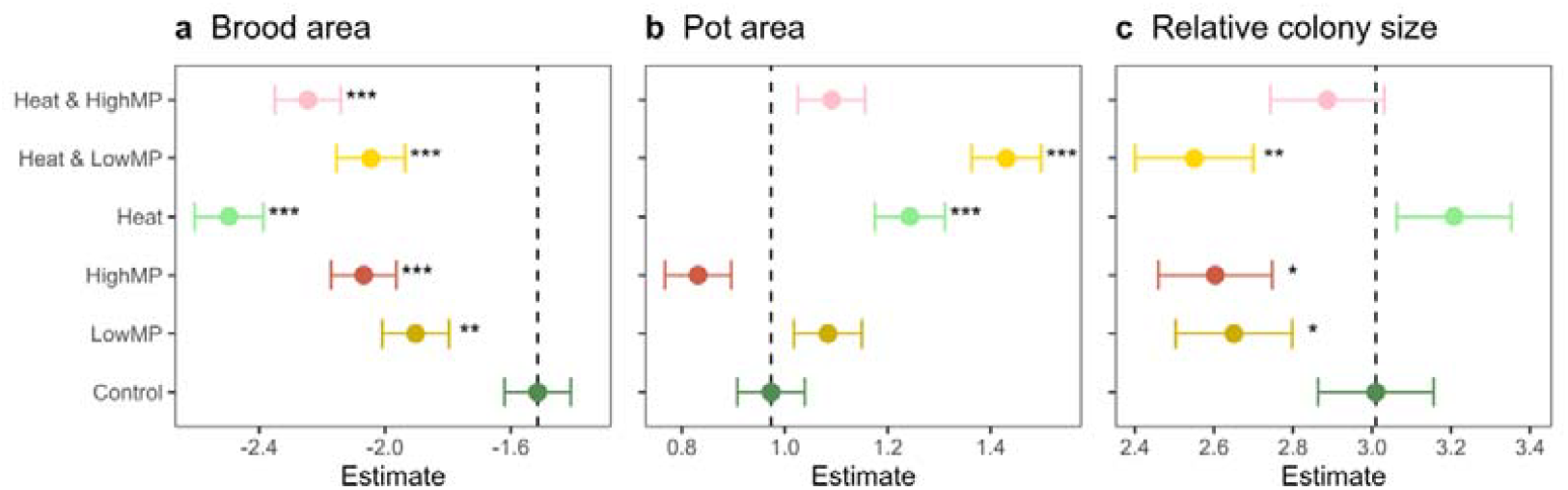
Treatment effects on colony level over time. Regression modelling results for **(a)** brood area, **(b)** pot area, and **(c)** relative colony size. Dependent variables were transformed according to formula. “***”: P-value < 0.001; “**”: P-value < 0.01; “*”: P-value < 0.05

The consistent significance across all variables under heat + low MP exposure—but not always under heat + high MP exposure—may stem from a threshold-dependent response, where low levels of plastic induce intensified physiological stress when compounded by thermal load, while high levels exceed the capacity for additional compensatory response. Thus, sublethal pollutant levels may have disproportionately high impacts under concurrent environmental stressors like climate change.

### Social level

We generated a series of social networks for each colony and calculated network densities (i.e., the general strength of connections) and the queen’s degree centrality rankings (i.e., the rank of the queen’s direct network connections relative to other bees in the network over time). The network density reflects the overall level of social interaction in the colony and whether the colony exhibits cohesive or fragmented social dynamics. The queen’s centrality ranking indicates her social integration and influence, and helps assess whether the queen plays a central or marginal role. Extended Data Fig. 11 shows the network series of a single colony across the chronic trial.

The social network densities of bumblebee colonies decreased with time (Extended Data Fig. 12a), suggesting less connectivity among individuals. Both MP and heat tended to stimulate the level of connectivity, and the combination of MP and heat induced stronger effects (p < 0.05 for low MP; p < 0.001 for other treatments) (Fig. 4a). The degree centrality of the queen also decreased with time, but its ranking was stable, showing a consistent relative status of the queen among her workers at different stages of colony (Extended Data Fig. 12b-c). Both MP exposure and heat showed significant positive effects on the queen’s centrality ranking (p < 0.01 for low MP or heat; P < 0.001 for other treatments) (Fig. 4b). Overall, social level effects of MP increased at least partially in combination with heat.

## Discussion

We investigated MP effects on bumblebees at individual, colony, and social level under optimal temperature and overheated conditions (for a summary of effect sizes see Fig. 1c). At individual level, MP exposure amplified activities related to foraging but not consistently stronger in combination with heat, while the opposite was true for nesting behavior that was mostly altered by combined treatments. At colony level, we found that MP exposure had a weak impact on colony size, but impaired brood development, especially in combination with heat. At social level, MP exposure stimulated network connectivity and the queen’s relative centrality, which was further exacerbated by heating. Under optimal temperature, MP effects are positively related with exposure level in general. However, under heated condition, low MP exposure induced some stronger effects than high exposure, for instance on pot area increase, nesting room activity, and queen centrality rank. This indicates that combined pressure of heat and low MP exposure likely falls within the hormesis range for bumblebee colonies (i.e., low doses of stress triggering beneficial adaptive responses) ^22,23^ with a biphasic pressure-response relationship.

Our findings suggest that MP pollution has complex, cross-level impacts on bumblebees and their pollination potential, which may be exacerbated with the increase of temperature. In general, the level of MP accumulation in our experiment is at the same order of magnitude (∼10^2^ items/mg) as reported in the literature^13^ (Supplementary Method 3), which is assumed a realistic exposure level of pollinators in anthropogenic environments. Temperature increase from 25 □to 30 □has led to increased nest abandoning^20^, which is aligned with our results that heat urged more passive bees in the foraging room and less in the nesting room. Moreover, bumblebees in the 30 □condition showed no significant difference in mortality and adult emergence^20^, which coincide with our findings that heat alone did not significantly affect the growth of relative colony size. At individual level, some studies have reported MP disturbance in food intake of honeybees. Balzani et al. found that honeybees consume more sucrose solution when exposed to low MP (0.5 mg/L)^16^, while other studies found decreased food intake with higher concentration (10 or 100 mg/L) of MP^13,24^. Dissections show that MP accumulate in the guts and damage tissue in the alimentary system^13,14,25^ and potentially obstruct mouthparts^16,24,26^. Therefore, the increased visitations on the sucrose solution feeder that we observed might be a compensating response to impaired alimentary systems and low feeding efficiency. Our findings at colony level are consistent with previous studies that have investigated MP effects on bee mortality and larvae exposure. Honeybees exposed to MP showed low mortalities^14,16,25^. However, foraging bees can transfer MP to the nest and to larvae^19^, thus threatening the colony offspring. This explains the additional brood loss and decreased growth in relative colony size in our findings. At the social level, effects of MP, especially in combination with heat, remain understudied. However, social connections can generally facilitate the spread of foraging knowledge within the colony^27^, and intense worker-queen interactions likely increases the dominance of the queen^28^. Our findings suggest that with MP and/or heat exposure, bumblebees communicate more actively especially with the queen in order to mitigate damages to the entire colony.

Our work provides a promising solution for advancing ecotoxicological assessment guidance within regulatory frameworks such as those of the Environmental Protection Agency (EPA)^29^ and the European Food Safety Authority (EFSA)^30^. This is particularly relevant given the recent growing attention to interactions of multiple pollinator stressors, such as climate change & land use^31^, or mixture of various agrochemicals^32^. By demonstrating the sublethal effects of MP and heating on pollinators at individual, social and colony level, our study underscores the need for long-term and multi-risk assessments in regulatory guidelines. Such assessments should aim to provide rich information on impacts on pollinator colony development and behaviors, which may profoundly alter the value of pollination service^33^. Future work should focus on expanding these studies to include a broader range of pollinator species and environmental contexts, as well as developing more advanced interdisciplinary tools to explore the impacts of multiple stressors. Additionally, collaborative efforts between researchers, policymakers, and industry stakeholders are essential to translate these findings into actionable policies towards the vision of the IPBES pollinator assessment report^7^ and the Post-2020 Global Biodiversity Framework targets.

## Methods

### Materials and devices

#### Bumblebee colonies acquisition and tagging

Our experiment was done on European bumblebees (*Bombus terrestris* Linnaeus, 1758), because the species is widely used as a major commercialized pollinator in agricultural production (such as tomatoes, melons, eggplants, etc.) globally^34^, and also a well-studied model organism for pollination studies. We ordered 30 queenright bumblebee colonies from BioBest®, with 38±4 adult bumblebees per colony and the entire nest structure (i.e., the wax pots and broods). We were unable to find a supplier to provide individual queens to start nest development in our research facilities. We were, however, assured that the BioBest® colonies were all started at the same time and under the same conditions. We then tagged individual bumblebee adults with mini matrix codes^21^, a miniature matrix code (3 mm × 3 mm) printed on water-proof paper that was then attached on the back of the bumblebee’s thorax by cyanoacrylate glue^21,35^ (Deli™) without affecting the mobility of the wings. To do this, a colony was N_2_-anaesthetized in a glove box, which is similarly effective to CO_2_-anesthesia^36^ but avoids side effects of hypercapnia^37^. We could not tag all individuals of all colonies when they were immobilized under the nest architecture and removing them would have damaged the nest structure. The nests were not fully removed at the beginning of the experiment, because bumblebee queens had made the initial investment to produce the nest and building a new colony would substantially disrupt the colony structure. After tagging, the entire colony, including the nest structure, was transferred into our behavioral monitoring system.

#### Customized bee monitoring system

We designed a customized behavioral monitoring system (Fig. 1a) for the experiments. Each monitoring system consisted of a wooden bee box and a movable camera system. The bee boxes had a nesting room (24 cm × 20 cm × 18 cm) and a foraging room (20 cm × 20 cm × 13 cm) connected with a tunnel (4 cm in diameter). The nesting room was covered with an opaque glass (near-infrared transparent) to create a dark environment within the visible light spectrum, which mimics the natural dark environment of bumblebee nesting. The foraging room was covered with a transparent glass lid. Walls and floors of both rooms were made of wood to avoid potential plastic pollution. Small wood grids (< 2 mm in width) on foraging room walls and nesting room floors allowed for ventilation. The floor of the foraging room had two holes (4 cm in diameter) as access to sucrose solution and pollen feeders. Bee boxes were supported by stand bars to prevent infection from the bottom and equipped with heating wires to adjust the temperature. To simulate natural lighting conditions, we placed full spectrum LEDs above the nesting rooms. We kept the colonies under 12h:12h L:D for an average of 20±1 days (the time variation resulted from the matrix code tagging applied to each colony). We used near-infrared LEDs (800 nm, F-Lighting™) to illuminate the bee boxes for the night and for the nesting room monitoring.

Our setup contained a total of 20 bee boxes with five bee boxes each placed together under one movable camera system to optimize the experimental setup for costs. Each camera system consisted of two programmable motion rails running in parallel. The slider on each rail moved simultaneously, carrying a monochrome industrial camera (2448 × 2048 pixels USB 3.0, MV-GE501GM-T, MindVision®), with one camera capturing images from the nesting room and another from the foraging room. The monitoring duration for each bee box was 5 minutes, resulting in a total monitoring cycle of 25 minutes for 5 boxes. Monitoring was continuously restarted throughout the entire experiment’s duration. To improve running speed and save storage space, we only saved detection data in csv format and discarded the images when the object detection algorithm was running. As a backup, a sample video was saved for each colony every 2.5 hours.

#### Feeding methods

Bumblebees were fed *ad libitum* and provided with sucrose solution (50% w/w) and dry pollen of oil-seed rape. The sucrose solution was placed in a glass container (Loikaw®), and then supplied by an automatic pump (NKP -DA, Kamoer®) through a feeding hole at the bottom of the foraging room. The glass container was refilled when empty (Extended Data Fig. 13) once a day. We covered the feeding hole with 2 mm × 2 mm metal grids to prevent bees from escaping. We provided dry pollen through another feeding hole in the foraging room and supply was also replenished once empty, every 4 days.

### Chronic trial design

#### Microplastic and temperature treatment

We used 30 μm polyethylene (PE) beads as MP pollutant as PE is the major component of mulching films and one of the main types of MP in agricultural landscapes^38^. It has also been found on honeybee samples globally^13,39^. We then assigned 30 bumblebee colonies to three different MP exposure levels – control (C), low (L), and high (H) respectively. The control groups were treated with negative 50% w/w sucrose solution, and low and high concentration groups contained sucrose solution of 10 mg/L MP and 100 mg/L MP, respectively. Dry pollens were mixed with MP of the same mass concentration (8×10^−5^ w/w). The regular refilling of sucrose solution and pollen allowed us to maintain a constant MP exposure level throughout the experiment (for validation of treatment effectiveness, see ‘Raman Spectroscopy’ section below). For each exposure level, we assigned colonies to normal (N, ∼25 □; the optimal temperature for bumblebees^40^), and warm (W, ∼30 □; realistic hot summer conditions in typical bumblebee habitats^41^) temperature conditions that were constantly monitored with data loggers in each bee box (A-BF™, BCL2024P, ±0.2□). Overall, we had six experimental groups (NC, NL, NH, WC, WL, WH) with five replicas each, randomly allocated to bee boxes in different rack locations.

#### Experimental setup

The experiment was set up in a dedicated room equipped to regulate temperature (±1°C) & humidity (70±10%) fully independent of the external environment. The number of required replicates (30 bee boxes) and available setup (20 bee boxes) required us to do the experiment in two subsequent iterations. The first experiment with 3 replicates (18 colonies) started on January 12^th^, 2024, and the second experiment with 2 replicates (12 colonies) started on June 5^th^, 2024. As the treatments were allocated balanced with all experimental conditions kept constant, induced variation from two experimental iterations is expected to be minimal (Extended Data Fig. 14 and Supplementary Method 4).

### Microplastic exposure validation

#### Sample preparation

To validate MP exposure success, we randomly collected bumblebee samples after both experimental iterations from all treatments. Following the experimental protocol^42^, 5 bumblebee samples per colony were pooled in 50 mL centrifuge tubes to reach sufficient quantities for Raman analysis (see next section). The samples were then freeze-dried to remove water, and their dry weights were recorded. Next, we randomly aggregated around 0.2 g bumblebee samples from each treatment. The aggregated samples were digested in 10 mL 30% H_2_O_2_ and 15 mL concentrated nitric acid for 48–72 hours at room temperature^13^ to eliminate organic matter from the bumblebee samples. Additional H_2_O_2_ and concentrated nitric acid were added dropwise to re-immerse particles adhering to the inner walls of the beakers. The mixture was then filtered through a 0.45 μm pore size polytetrafluoroethylene (PTFE) membrane filter (50 mm in diameter, Devstlab®).

#### Raman Spectroscopy

Raman spectroscopy was used to confirm accumulated PE MP in bumblebees. The Raman spectra were recorded in the air using a WITec confocal Raman microscope (Alpha 300R, Germany) equipped with a 532 nm solid-state laser (≥ 75 mW). The collected Stokes Raman was dispersed by a UHTS 300 mm spectrometer and detected using an electron-magnified charge-coupled device (EMCCD) thermoelectrically cooled to − 60 °C. All single spectra were achieved under a 10× long-focus objective lens (Zeiss EC Epiplan) at room temperature over the wave number range of 0 – 4,000 cm^−1^ (600 gratings/mm) with an integration time of 1 s and 10 accumulations. To account for possible heterogeneity in residue distribution across the filter membrane, we randomly selected 6 field areas in each sample and recorded Raman spectra for suspected PE MP. Spectra of standard PE particles (the same product used for chronic exposure to bumblebees) were also recorded. We removed intrinsic autofluorescence backgrounds^43^, performed Savitzky-Golay smoothing (Extended Data Fig. 1), and then identified MP of the same type by principal component analysis and K-means clustering (Extended Data Fig. 2a-b and Supplementary Method 1). MP accumulation (particles per mg) is calculated as

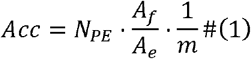

where *N*_*PE*_ is the number of identified PE MP, *A*_*f*_ is the total area of the filter, *A*_*e*_ is the examined area, and *m* is the sample weight.

### Computer-vision-based monitoring

We used object detection models based on MMDetection^44^ to identify target objects from images captured by our monitoring cameras. The object detection model is able to identify four different categories of objects (Fig. 1b), i.e., the 5×5 mini matrix codes (BEEtags ^21^), bees (individual adult bumblebees), pots (wax pots in the nest, used for food storage), and broods (enclosed lumps that contain eggs, larvae or pupae). The tags detected by the model were then decoded according to Crall’s principles^21^ (i.e., a 5×3 area of binary identification codes plus a 5×2 area of error check zone). We firstly used 31 images with 3101 labeled objects and trained a preliminary object detection model with an F1 score of 0.7820 (Extended Table 1 and Extended Data Fig. 15) and used it for the chronic trial in January 2024. To improve the detection reliability of the broods (recall = 0.4079), we performed more training epochs using 59 additional images with 5780 labeled objects from the first trail. The new model (Extended Data Table 2 and Extended Data Fig. 16) has an F1 score of 0.8326 and showed better performance especially on brood detections (recall = 0.6785). The new model was used for the second trail in June 2024. Moreover, as brood and pot area does not change much within 2.5h time steps, we used the new model to improve data quality for brood and pot area detection of the first trial based on the backup videos recorded every 2.5h (for validation see Supplementary Method 5 and Extended Data Fig. 17).

### Index design and statistical analysis

#### Individual behavior indices

At the individual level, we measured the activity, labor division, and intraday variance of bumblebee colonies. We classified the labor types of each individual according to their spatial locations (Extended Data Table 3 and Extended Data Fig. 18). For each type of labor, we calculated the proportion of assigned bumblebees among the total number of simultaneous bumblebee detections. As the proportion data ranges from 0 to 1, we used a logit link function to transform the original data, and then calculated the difference between each day and the first day of the trial:

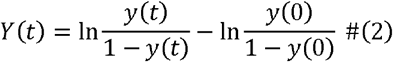

where *y*(*t*) is the original data (pot area or brood area) at time *t*; *y*(0) is the original data of the first day; and *Y*(*t*) is the transformed data that’s used as dependent variable for linear regression. We constructed the linear regression model as

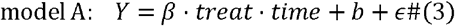

where *Y* is the dependent variable; *treat* is the categorical variable indicating the treatment to the colony; *time* is the scaled time variable (*time = day/*20); *β* is the coefficient that represents time effect of the treatment; *b* is the intercept; and *ϵ* is the residual item. We then tested the differences between coefficient estimates of treated groups and the control group (Supplementary Method 6).

We used average moving speeds (measured in pixels per second) to represent the level of activity. Since speed calculation relies on individual tracking across frames, we only measured activity of the tagged bumblebees, i.e., the queen and early emerged workers. We calculated the difference between each day’s value and the first-day value of each colony, *Y*(*t*) = *y*(*t*) − *y*(0), and then used model A to analyze the impact on activity changes and on labor division changes. For the latter, the full dataset including tagged and untagged bees were used, because determining the proportions of labor types does not require the individuals to be uniquely identifiable.

#### Colony level indices

At colony level, we measured nest structures and population growth. We described the structure of the nest by calculating the proportion of brood area and nectar pot area in the image, with values for both areas ranging from 0 to 1.

Colony size was estimated as the maximum number of simultaneous bumblebee detections every 12 hours. Colony sizes plateaued after around 12 days (Extended Data Fig. 9c) indicating that colonies have approached the maximum potential population. We then used the maximum value of each colony to scale the colony size data into 0 – 1 as ‘relative colony size’. Then we to transformed relative colony size data according to formula (2) and applied regression model A to analyze the impact of MP and/or heat on colony size changes. To investigate the impact on maximum colony size, we used the maximum value of each colony as dependent variable and constructed a second regression model as

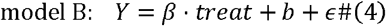

#### Social level indices

Network analysis has been widely applied in social sciences and lately ecology to study structures and the links among individuals^45,46^. Uniquely identifiable tracing of individuals in our bumblebee colonies allows us to apply network analysis tools as well. Ideally, in a fully presented network, each node would represent one traceable individual (Extended Data Fig. 19a); however, our experiment included interactions between both tagged and untagged individuals. Therefore, we visualized a bumblebee social network as a queen node, some tagged worker nodes, an aggregated node of untagged individuals, and the connections (edges) between them (Extended Data Fig. 19b). Considering the grey box characteristic of the aggregated node, we modified traditional ways to calculate network properties accordingly.

In the following mathematical deductions, we started with the traditional definitions of network properties based on a fully presented network (each node represents an individual) and ended with formulas that only contain known quantities.

We used weighted undirected graphs to represent bumblebee social networks and define the weight of the edge that connects node *i* and *j* as the probability of physical contact between individual *i* and *j*. Therefore, the weight is calculated as *w*_*ij*_ = *E*_*ij*_/*F*_*ij*_, where *E*_*ij*_ is the number of physical contact events; *F*_*ij*_ is the number of frames where both *i* and *j* are detected. Here, a “physical contact event” is defined as when the object detection bounding boxes of two individuals are overlapping (Extended Data Fig. 20). It can be inferred that *w*_*ij*_ ranges between 0 and 1 and bigger weights indicated stronger connections. When two nodes are disconnected, the “edge” between them is mathematically defined as 0. A network is generated every 2 days for each colony to obtain enough data for network construction.

#### Weighted network density

We define the weighted density of a bumblebee social network as

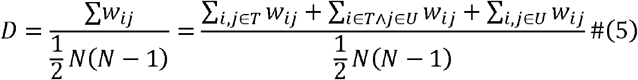

where *w*_*ij*_ is the weight of the edge that connects *i* and *j*; *N* is the population of the bumblebee colony (number of nodes); *T* is the set of tagged individuals; *U* is the set of untagged individuals. The first item in the numerator is known. And the second item can be converted into

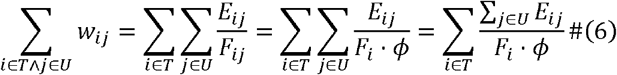

where *F*_*i*_ is the number of frames with the tagged individual *i*; *ϕ* is the possibility that untagged individual *j* is detected. We use the product of *F*_*i*_ and *ϕ* to represent *F*_*ij*_ because the detection of *i* and *j* should be considered as independent. We assume that the detection possibilities for all untagged bumblebees are the same, thus can be obtained from Extended Data Table 1 and 2 as *ϕ* = 0.8018 for the preliminary model and *ϕ* = 0.8723 for the final model. Σ_*j*∈*U*_ *E*_*ij*_ can be calculated by summing up all physical contact events between tagged individual *i* and any untagged individual *j*. The third item in the numerator can be converted into

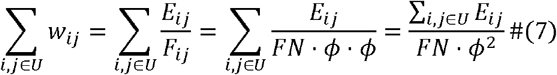

where *FN* is the total number of frames. Σ_*i,j*∈*U*_ *E*_*ij*_ can be calculated by summing up all physical contact events between untagged individuals. Finally, we calculate the weighted density *D* of a bumblebee social network as

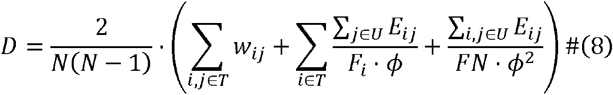

We transformed network density data with formula (2), and then used model *A* to analyze the impact on network densities.

#### Weighted degree centrality of the queen node

We defined the weighted degree centrality of the queen node as

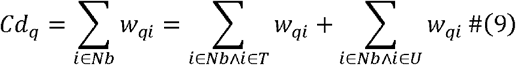

where *Nb* is the set of neighboring nodes to the queen node *q*. The second item in the formula can be converted into

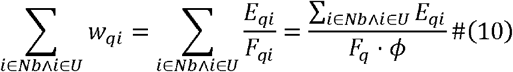

where Σ_*i*∈*Nb*Λ*i*∈*U*_ *E*_*qi*_ can also be directly measured. Therefore, the degree centrality of the queen node can be expressed as

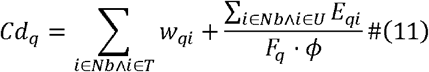

To assess the queen’s relative status within the colony, we take the centrality ranking of the queen among all nodes as the dependent variable. We calculated the difference between each period’s ranking and the ranking in the first network, *Y*(*t*) = *y*(*t*) − *y*(0), and then used model A to analyze the impact on queen centrality ranking. Raman spectra data analysis were performed in Matlab (version R2020b), and all other data handling and statistical analysis in this work were performed in R (version 4.3.3). The following R packages were used: ‘gglpot2’ and ‘ggpubr’ for visualization; ‘car’, ‘lmerTest’ and ‘AICcmodavg’ for data cleaning and analysis, ‘igraph’, ‘ggraph’, ‘tidygraph’ for network analysis.

## Data availability

The data that support the findings of this study are available from the corresponding author upon reasonable request.

## Acknowledgements

We would like to thank Dr. Lishun Wang and Dr. Xin Yuan for their generous help in the object detection model framework. We thank Dr. Dimitrios Karanikolopoulos for his support in matrix code tracking techniques at the early stage of the experiment. We thank Dr. Acheampong Atta-Boateng for his insightful suggestions on social network analysis.

## Author contributions

D.S., S.J. and T.C.W. conceived and planned the study. D.S. and S.J. designed the beekeeping system. D.S. and M.B. developed the monitoring system. D.S. trained the object detection model. S.J. and T.C.W. prepared the controlled bee room. D.S., S.J. and M.B. build up the experiment equipment. D.S. and S.J. prepared bumblebee colonies, sucrose solution and pollen. D.S. and S.J. conducted the Ramen spectroscopy detection. D.S. analyzed the experiment data. D.S, S.J. and T.C.W. wrote the main text. D.S. wrote the supplementary materials. All authors reviewed and edited the text.

