## Extended Data for "Long-term exposure to microplastics and heat affects bumblebee behavior patterns, colony development and social networks"

† Contributed equally

### Extended Tables

**Extended Table 1 | Preliminary object detection model evaluation.** Score threshold = 0.7, IoU threshold = 0.5.

| Category | Precision | Recall |
| --- | --- | --- |
| “tag” | 0.9241 | 0.6577 |
| “bee” | 0.9751 | 0.8018 |
| “pot” | 0.9641 | 0.7902 |
| “brood” | 0.9375 | 0.4079 |
| Average | 0.9502 | 0.6644 |
| F1 score | 0.7820 | |

**Extended Table 2 | Final object detection model evaluation.** Score threshold = 0.7, IoU threshold = 0.5.

| Category | Precision | Recall |
| --- | --- | --- |
| “tag” | 0.9902 | 0.8145 |
| “bee” | 0.9210 | 0.8723 |
| “pot” | 0.9428 | 0.8564 |
| “brood” | 0.8991 | 0.7481 |
| Average | 0.9383 | 0.8228 |
| F1 score | 0.8768 | |

**Extended Table 3 | Definition of different labor types.**

| Labor type | Definition |
| --- | --- |
| Pollen | Bounding box of the bee has overlap with the pollen feeder. |
| Sucrose | Bounding box of the bee has overlap with the sucrose feeder. |
| Passive foraging room | Bee is in the foraging room but has no overlap with feeders. |
| Nursing | Bee has overlap with the nest architecture (brood and pot area) |
| Passive nesting room | Bee is in the nesting room but has no overlap with the nest architecture. |

### Extended Figures

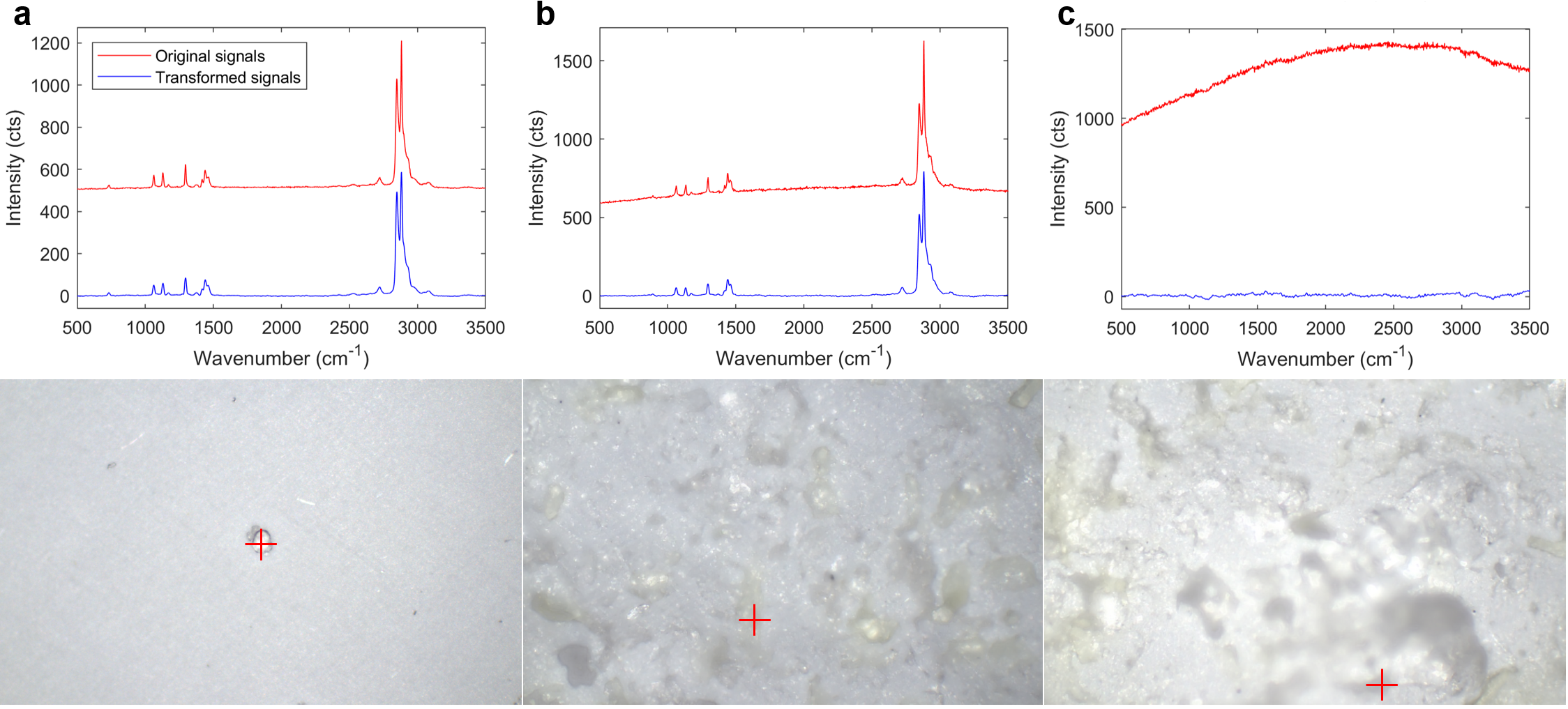

Extended Data Fig. 1 | Raman spectra and microscopic images (10× magnification) of microplastic particles. (a) Standard PE microplastic particles, (b) PE particles detected in bumble bee samples, and (c) uncertain particles from bumblebee samples. The red cross marks the particle of interest. Red curves are the raw signals, and blue curves are the transformed signals after autofluorescence backgrounds removal and Savitzky-Golay smoothing.

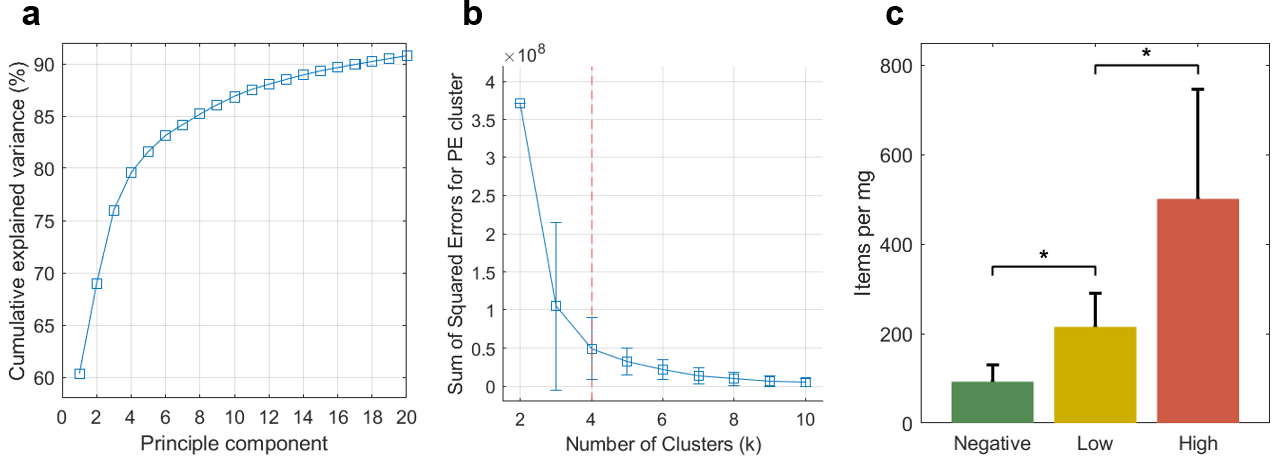

Extended Data Fig. 2 | Identifying potential PE microplastics. (a) Cumulative explained variance of the principal components. The top 18 principal components accounted for the majority (>90%) of the variance in the dataset, thus were chosen to conduct K-means clustering. (b) Sum of squared errors (SSE) for the known PE cluster after 1000 iterations of K-means clustering. The elbow point is set at 4 because this is the where the SSE curve starts to flatten out, meaning further increase of the number of clusters contributes little to clustering quality. (c) Microplastic accumulation in bumblebee samples (number of PE microplastics per mg) determined by equation (1) in the main text. “*”: P-value < 0.05 (T-test, n=6 in each group). For further details see Supplementary Method 1.

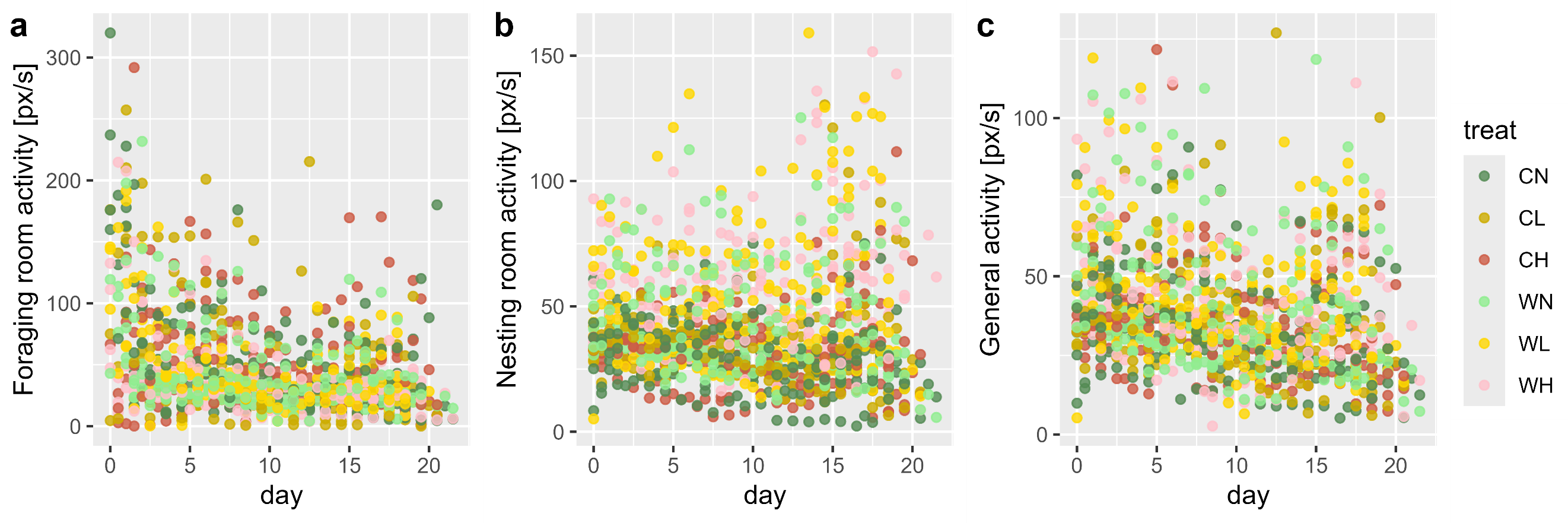

Extended Data Fig. 3 | Time series of activity (i.e., average moving speed of tagged individuals) within (a) foraging rooms and (b) nesting rooms, and (c) of the entire colonies.

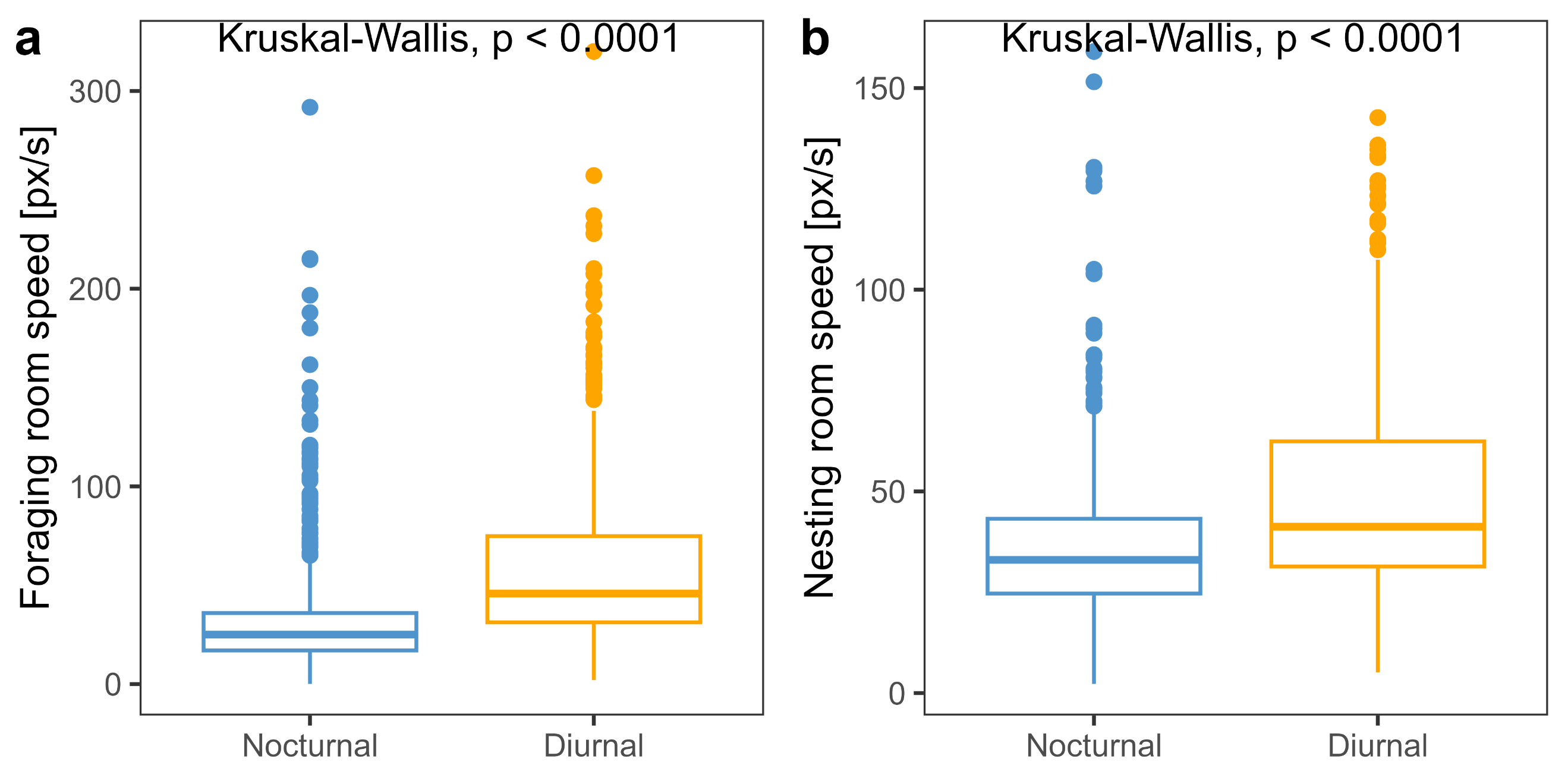

Extended Data Fig. 4 | Activeness (moving speed) difference between diurnal times and nocturnal times. Kruskal-Wallis tests are performed between diurnal data and nocturnal data, and P-values are provided.

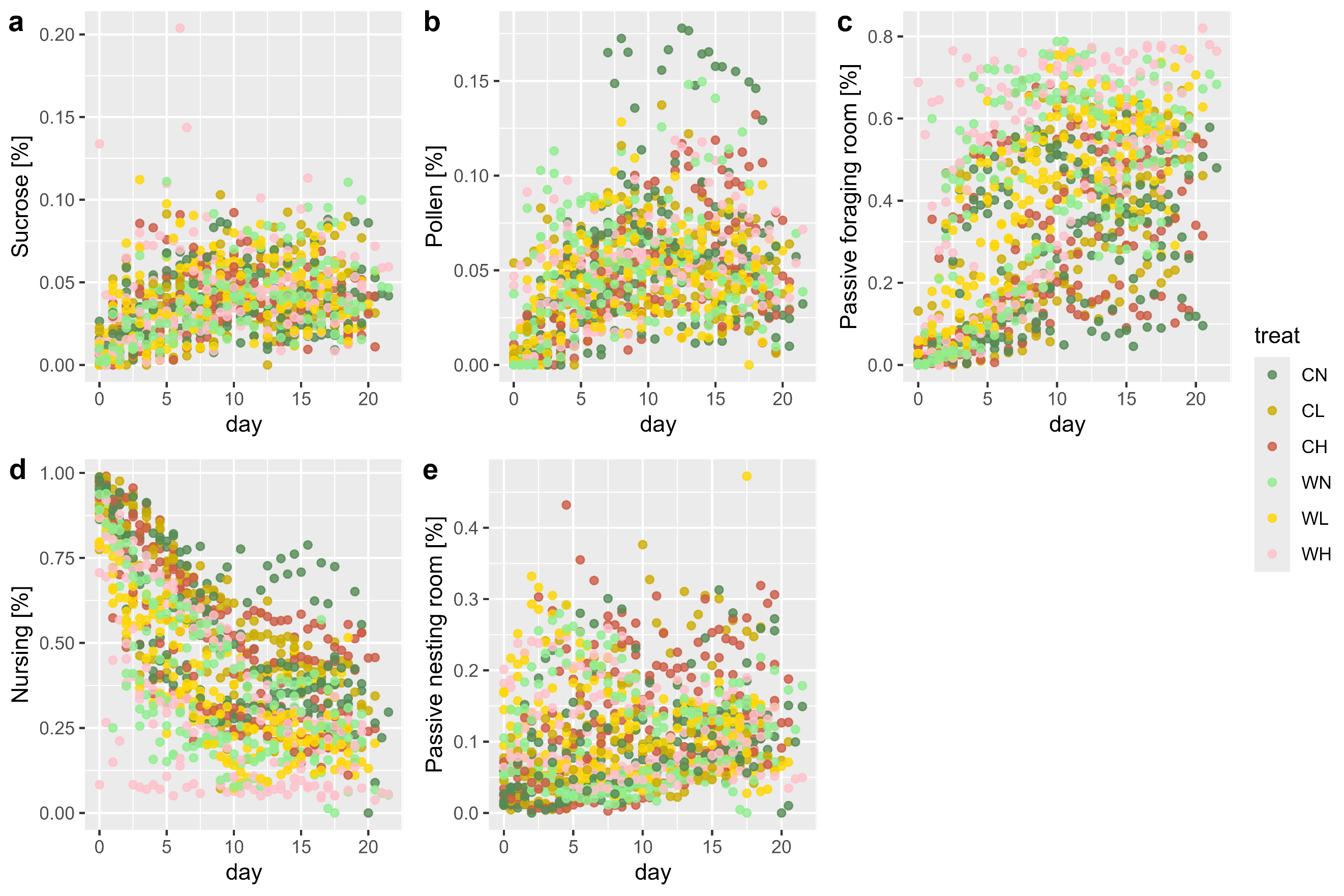

Extended Data Fig. 5 | Time series of the proportion of bees assigned to different labor types, namely (a) “out”, b) “^2–4^sucrose”, c) “pollen”, d) “nursing” and e) “rest”. For definitions of labor types, see table S2.

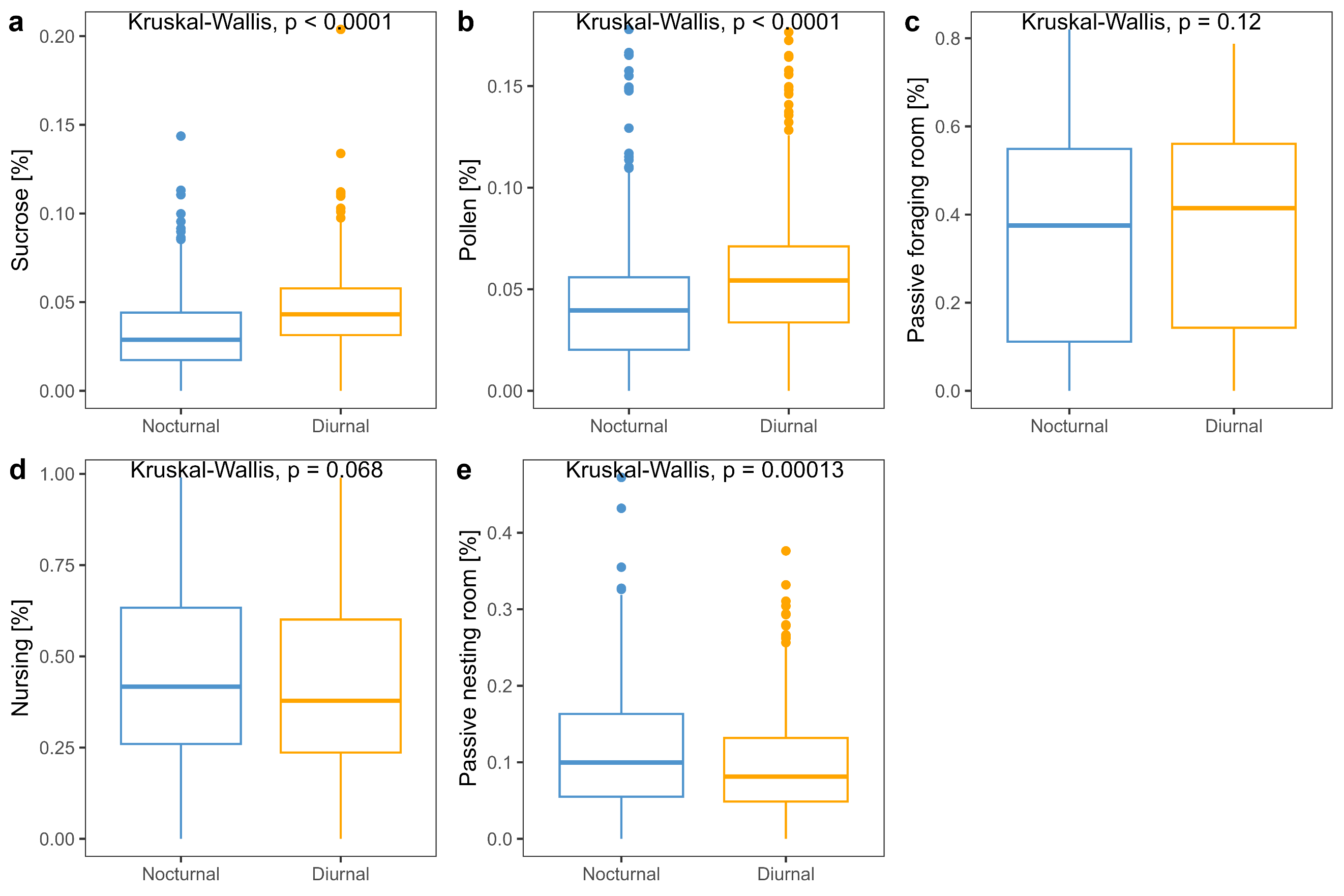

Extended Data Fig. 6 | Labor division difference between diurnal times and nocturnal times. Kruskal-Wallis tests are performed between diurnal data and nocturnal data, and P-values are provided.

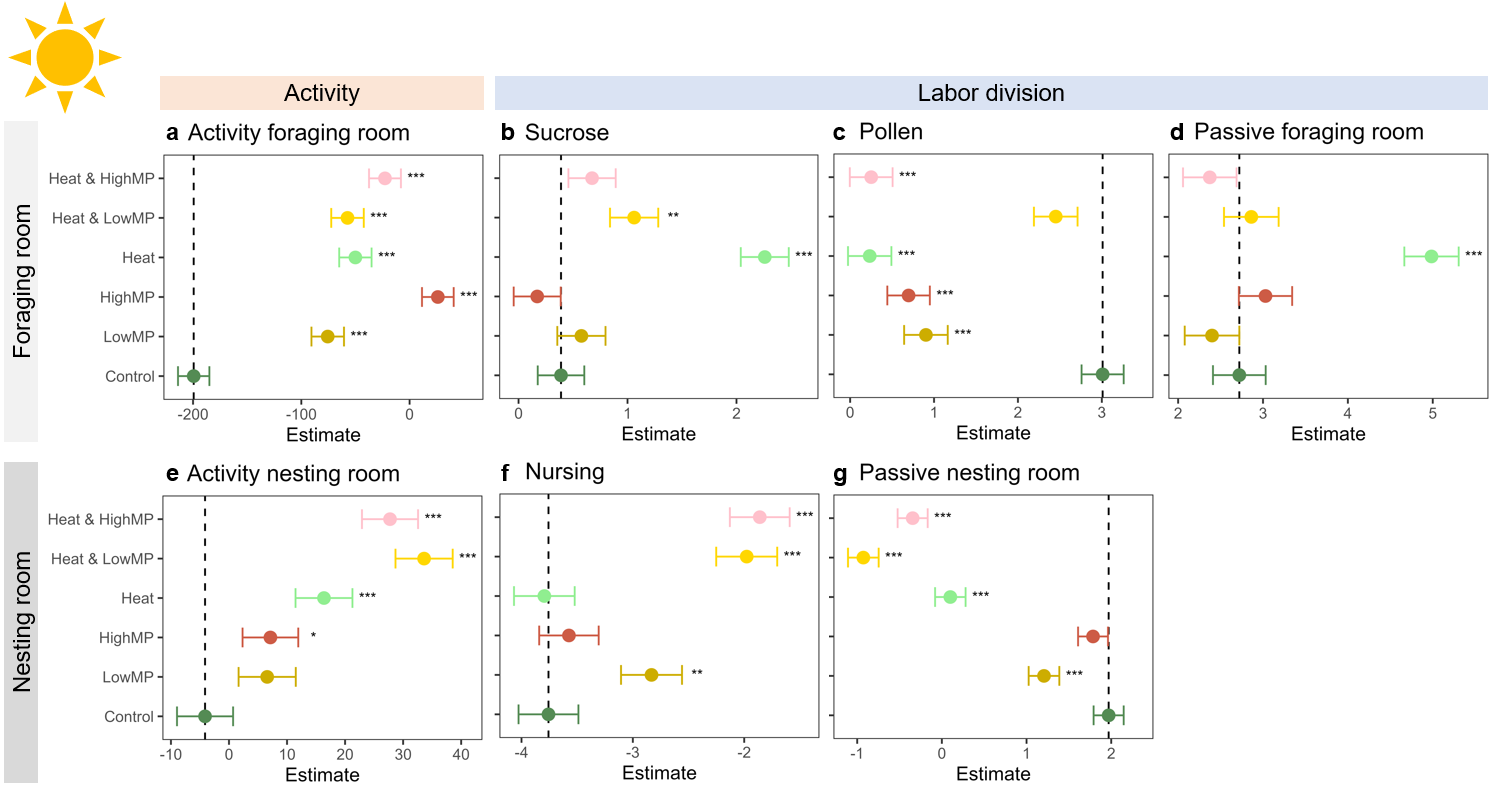

Extended Data Fig. 7 | Diurnal (6:00 – 18:00) individual level estimates of time effects using model A. Panel (a) is individual activity in the foraging room, and (b), (c), and (d) are to foraging on sucrose solution, foraging on pollen, and passive behaviors in the foraging room. Panel (e) is individual activity in the nesting room, and (f) and (g) are nursing behaviors and passive behaviors in the nesting room. Labor division variables [Panel (b) - (d), (f) - (g)] were transformed according to formula $\left( \boldsymbol{1} \right)$. For definitions of labor division, see Extended Data Table 3. “***”: P-value < 0.001; “**”: P-value < 0.01; “*”: P-value < 0.05.

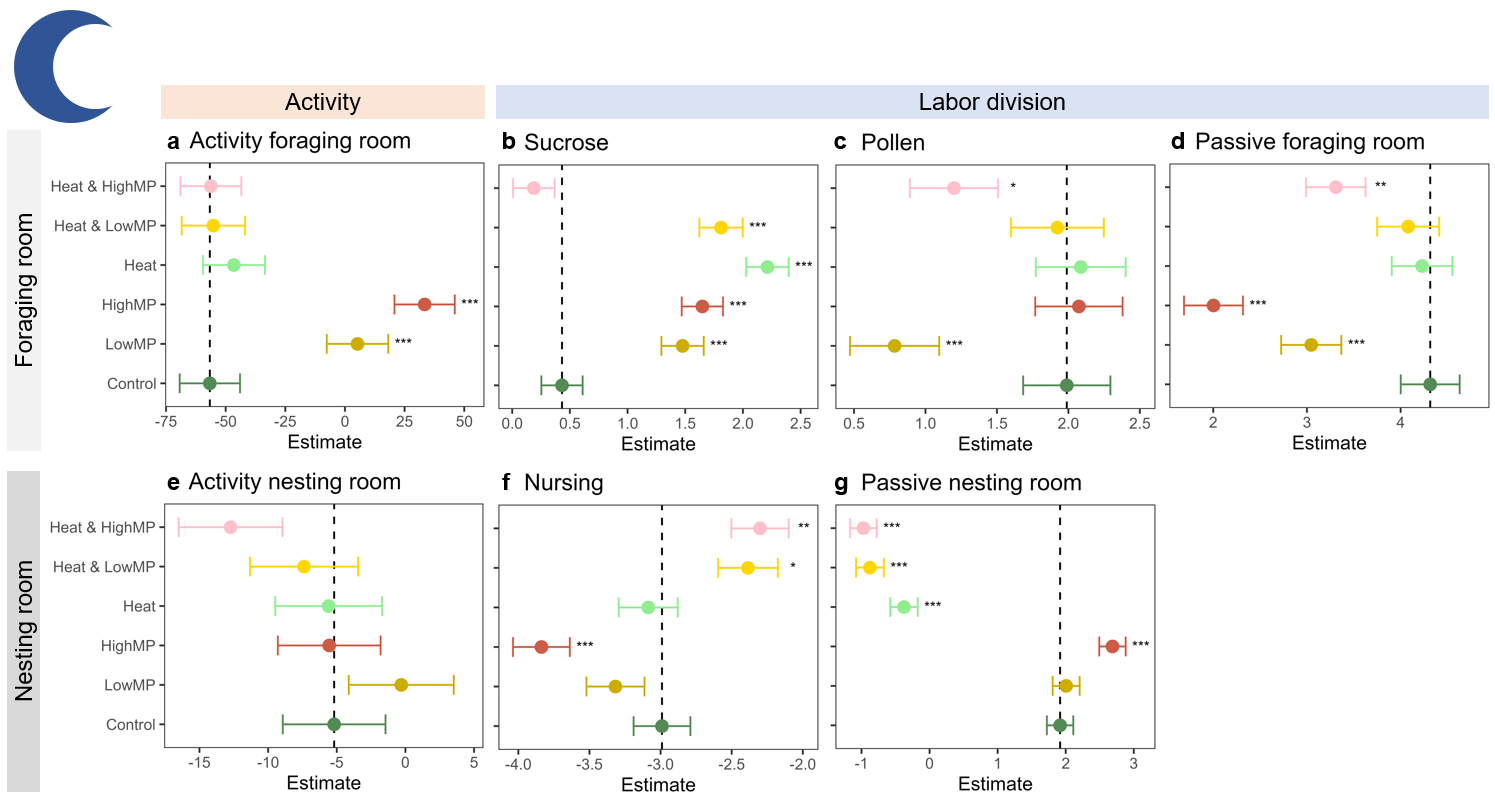

Extended Data Fig. 8 | Nocturnal (18:00 – 18:00) individual level estimates of time effects using model A. Panel (a) is individual activity in the foraging room, and (b), (c), and (d) are to foraging on sucrose solution, foraging on pollen, and passive behaviors in the foraging room. Panel (e) is individual activity in the nesting room, and (f) and (g) are nursing behaviors and passive behaviors in the nesting room. Labor division variables [Panel (b) - (d), (f) - (g)] were transformed according to formula $\left( \boldsymbol{1} \right)$. For definitions of labor division, see Extended Data Table 3. “***”: P-value < 0.001; “**”: P-value < 0.01; “*”: P-value < 0.05.

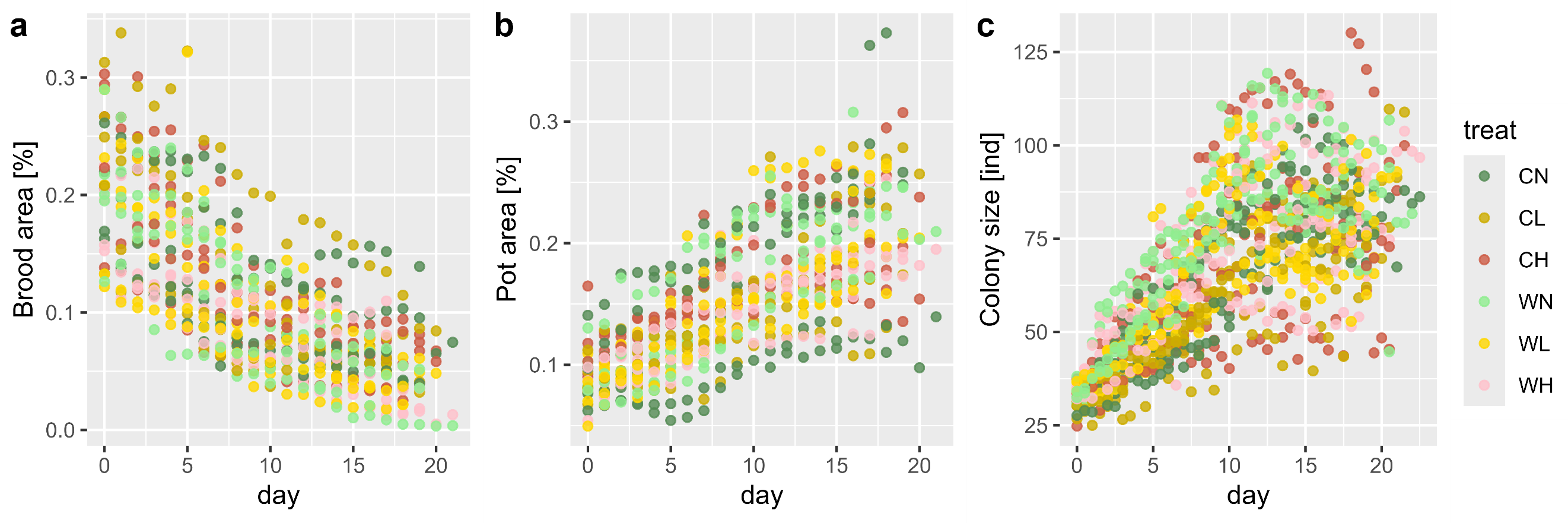

Extended Data Fig. 9 | Time series plot of colony level indices. Panel (a) is the brood area as a proportion of the entire image, (b) is the pot area as a proportion of the entire image area, and (c) is the relative colony size as a proportion of the maximum population of each colony.

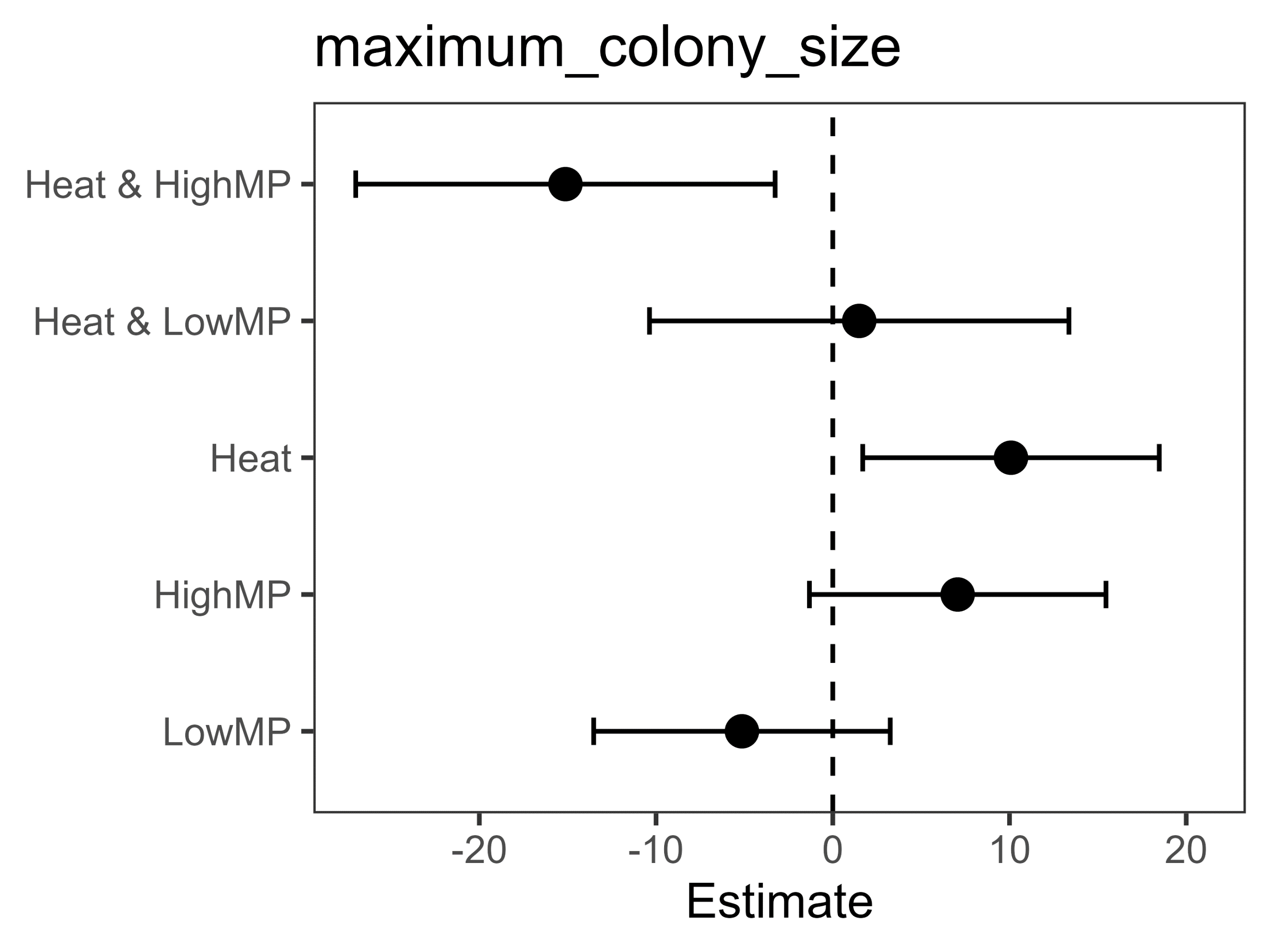

Extended Data Fig. 10 | Estimates of microplastic and/or heat effects on maximum colony size (model B). “***”: P-value < 0.001; “**”: P-value < 0.01; “*”: P-value < 0.05.

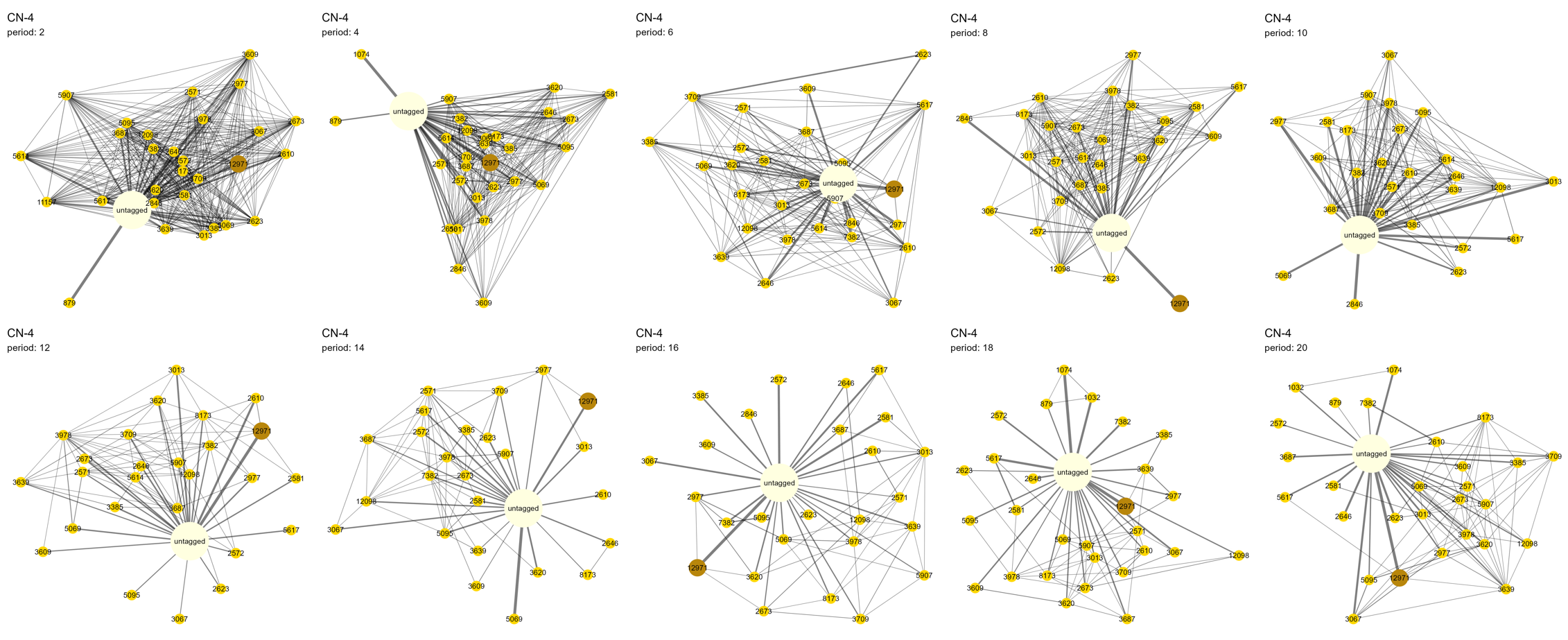

Extended Data Fig. 11 | Bumblebee social network time series, using colony CN-4 as an example. Orange nodes represent tagged workers, brown node represents the queen, and pale-yellow node represents the aggregate of all untagged bees. Nodes are arranged according to Davidson and Harel’s simulated annealing algorithm.

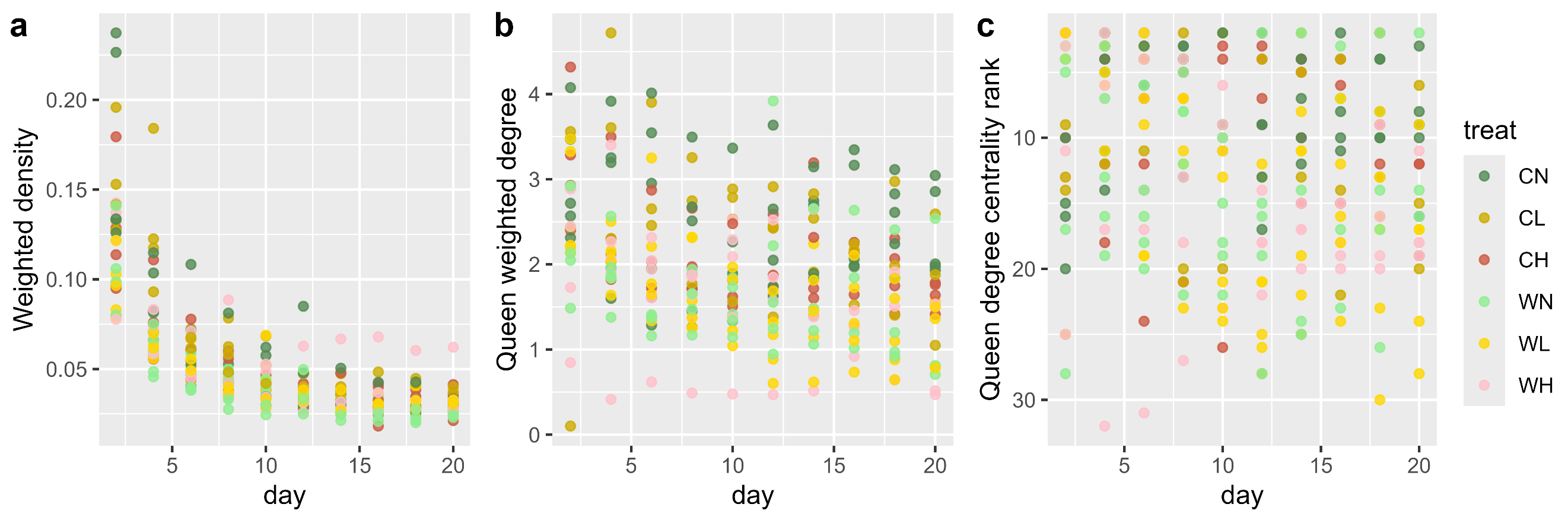

Extended Data Fig. 12 | Time series of network properties. Shown indices are (a) network weighted density, (b) queen weighted degree centrality; and (c) queen centrality ranking.

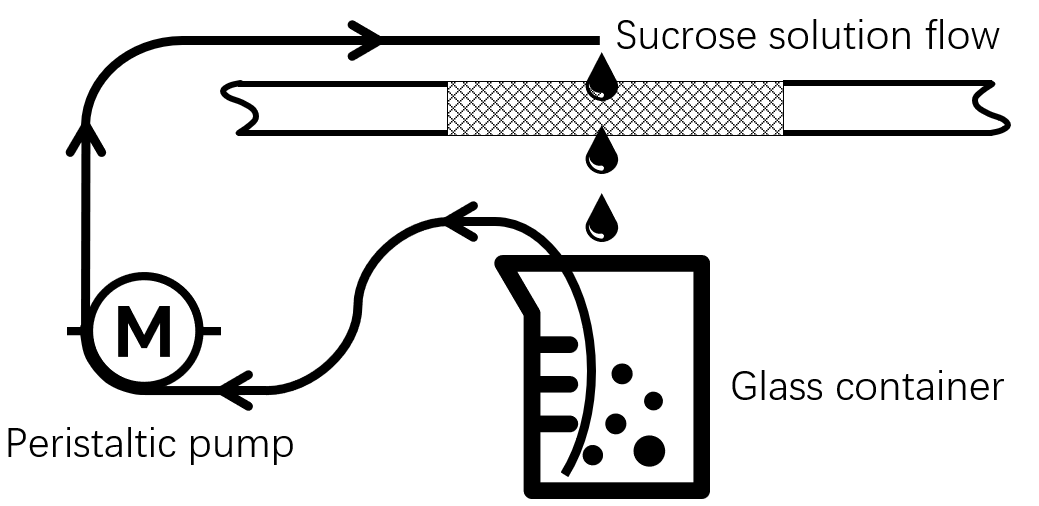

Extended Data Fig. 13 | Feeding method of sucrose solution. A peristaltic pump with a silicone tube is used to pump sucrose solution from the glass container to the inside of the foraging room. The sucrose solution then drops back into the glass container through the grids, if not foraged by bumblebees.

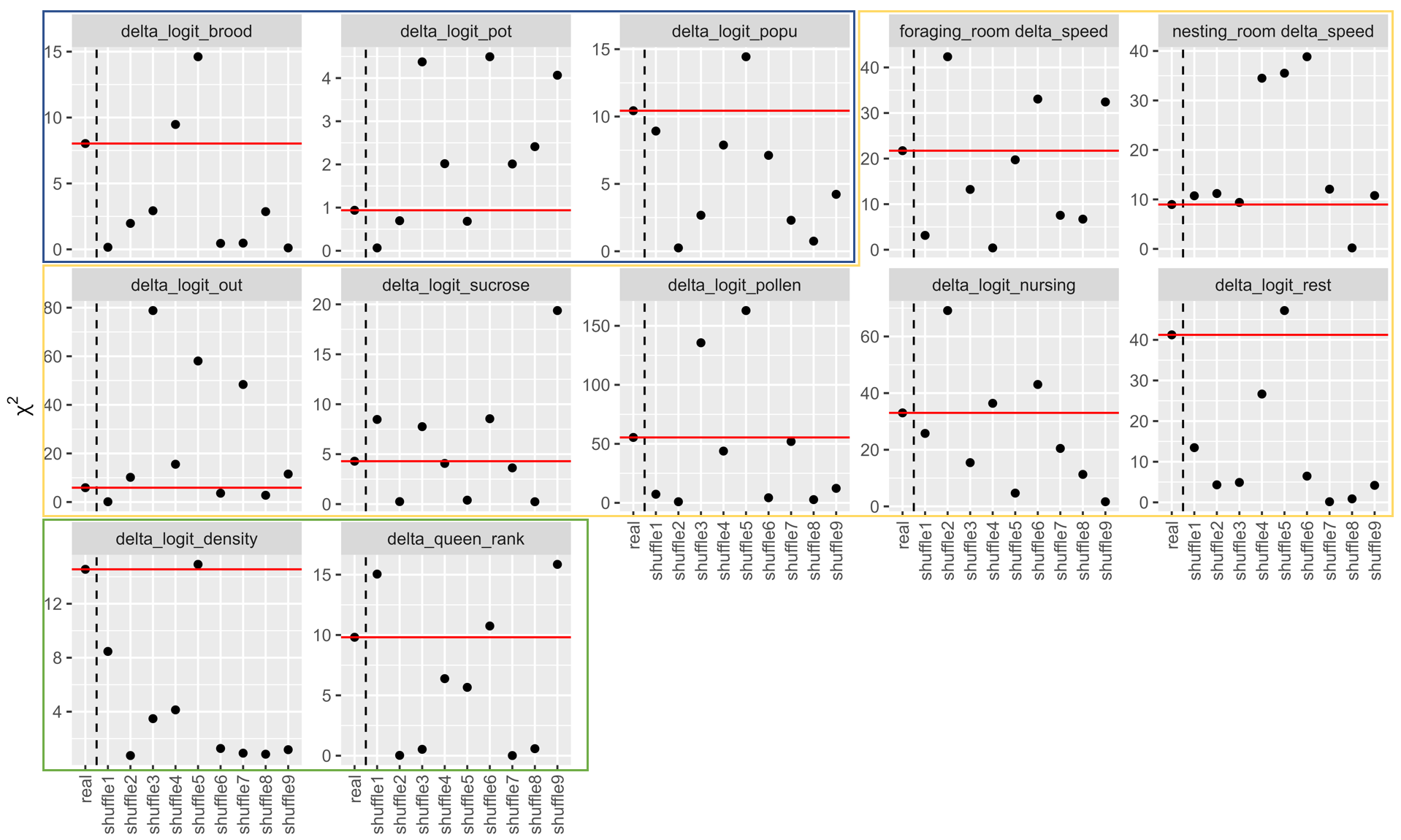

Extended Data Fig. 14 | Kruskal-Wallis test results ($\boldsymbol{\chi}^{\boldsymbol{2}}\boldsymbol{)}$ for two experimental iterations and shuffled combinations. Circled in blue, yellow and green are dependent variables at colony level, individual level and social level respectively. Red horizontal lines mark the $\boldsymbol{\chi}^{\boldsymbol{2}}$ level of the real categorization of experimental iterations, which fell between the maximum and minimum values of shuffled combinations, meaning no additional variation induced by experimental iterations.

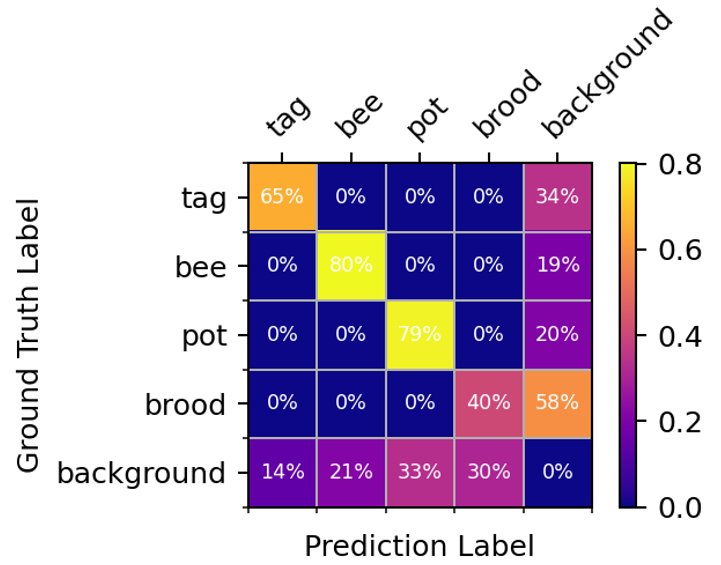

Extended Data Fig. 15 | Normalized confusion matrix of the preliminary object detection model. Score threshold = 0.7, IoU threshold = 0.5.

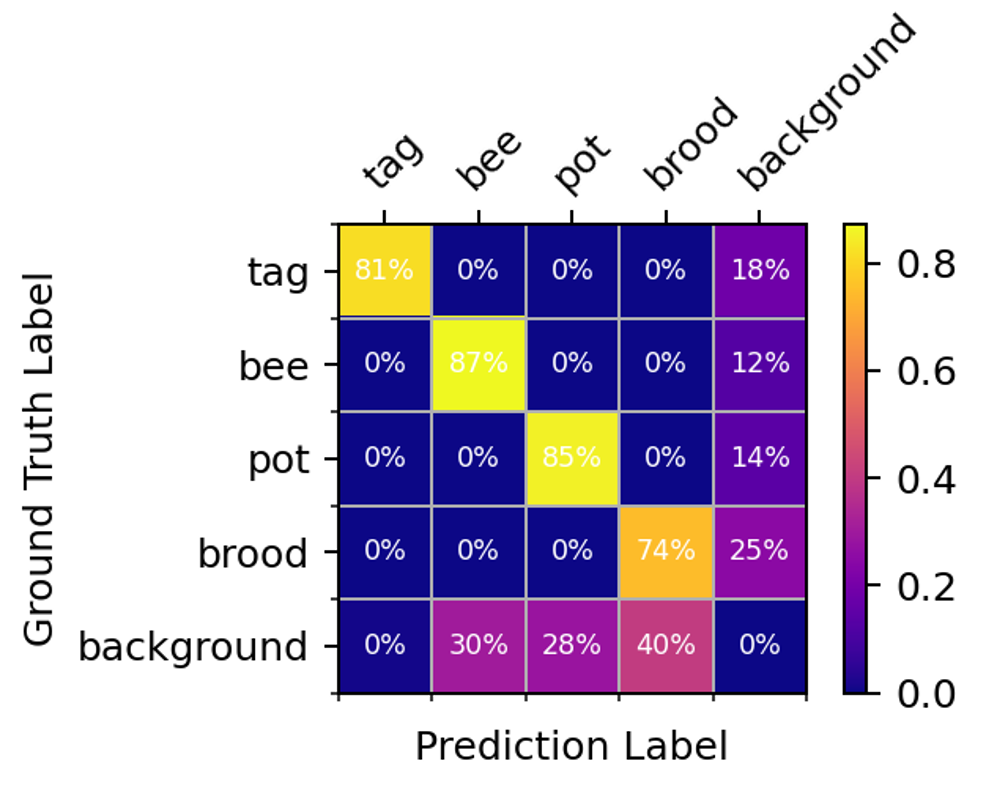

Extended Data Fig. 16 | Normalized confusion matrix of the final object detection model. Score threshold = 0.7, IoU threshold = 0.5.

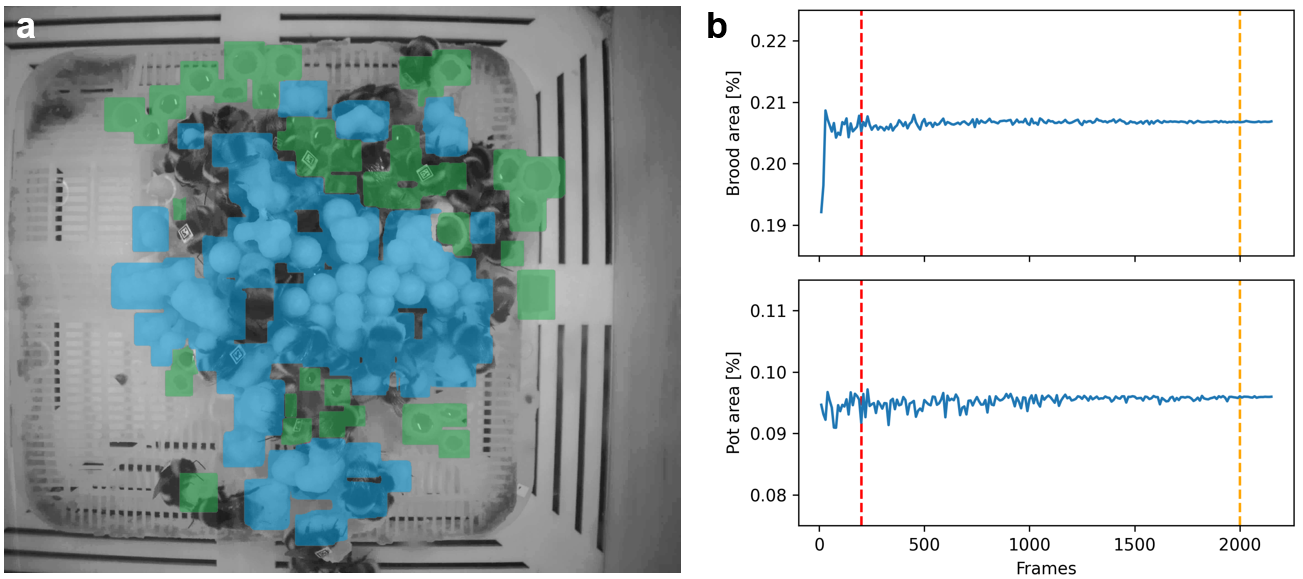

Extended Data Fig. 17 | Detection of brood area and pot area with sample videos. (a) The nest structure of an example colony, with brood area shaded blue and pot area shaded green. (b) Detected brood area and pot area against the accumulation of sampling frames. Red dashed line marks the approximate number of frames with one sample video, and orange dashed line marks the number of sample video frames after 24 hours. The curves stabilized before the number of frames reaches the orange dashed line, meaning sample videos were sufficient to reconstruct the nest structures on a daily basis.

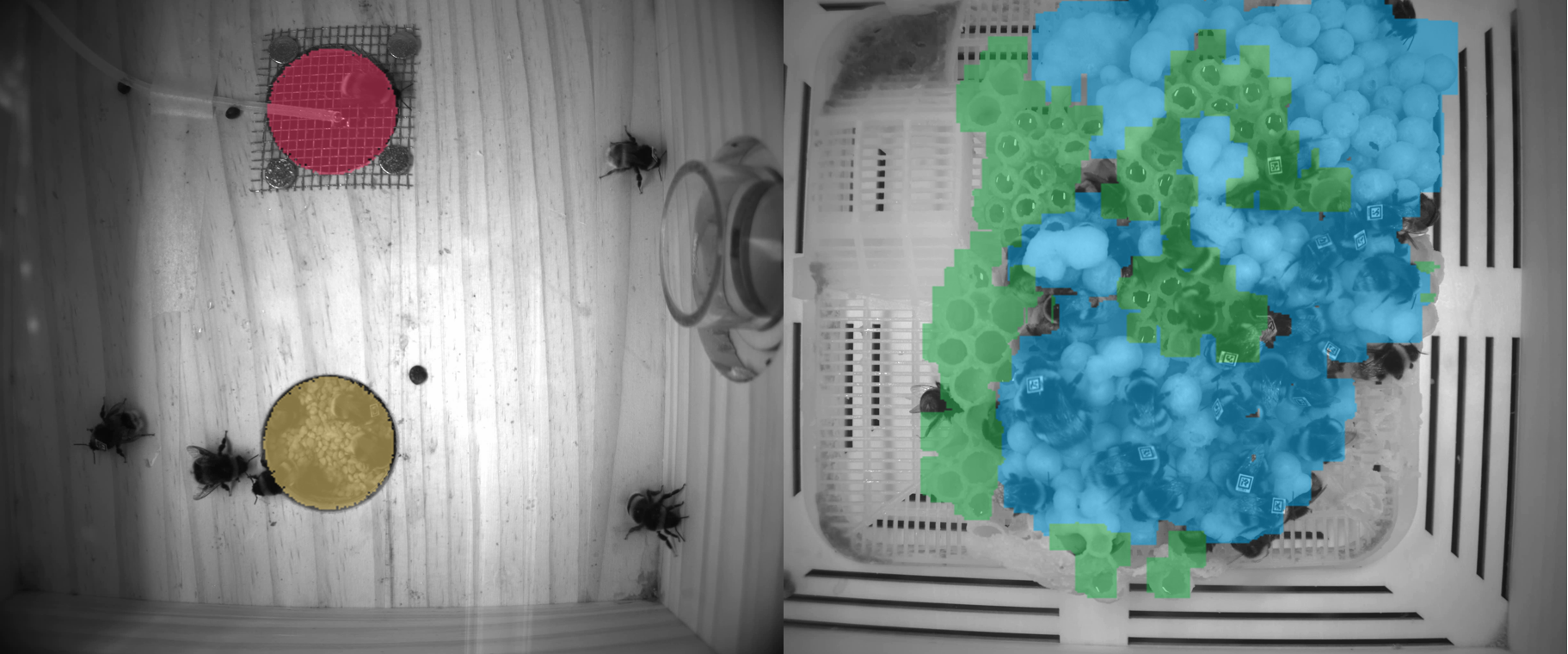

Extended Data Fig. 18 | Labor types of individuals are determined according to their spatial locations. Sucrose feeder and pollen feeder are shaded in red and yellow in the foraging room. Brood area and pot area are shaded in blue and green in the nesting room.

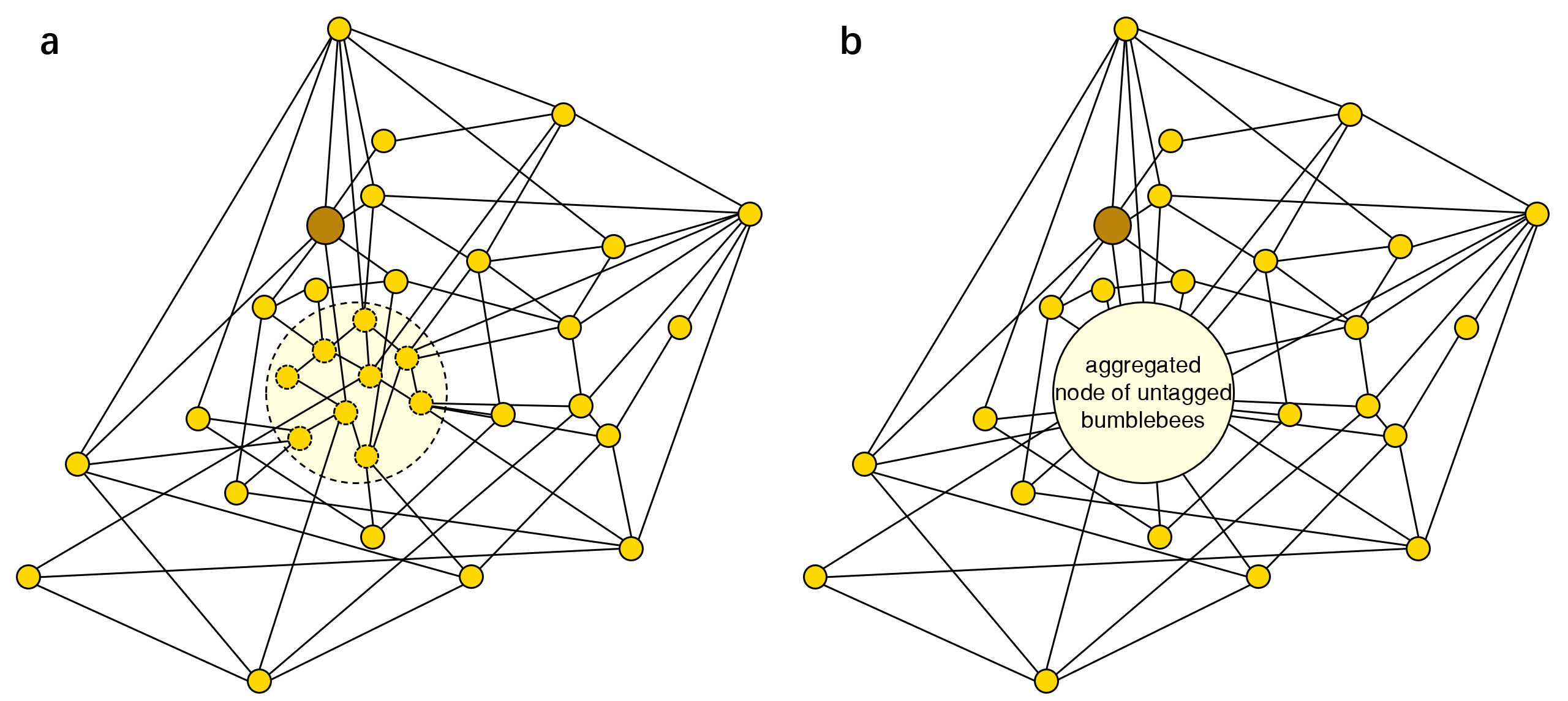

Extended Data Fig. 19 | Social network of a bumblebee colony based on physical interactions. Golden nodes represent workers, and the brown node represent the queen. The shaded area in (a) includes untagged bumblebees that were untraceable. These untagged individuals were aggregated as one node in pale yellow in (b).

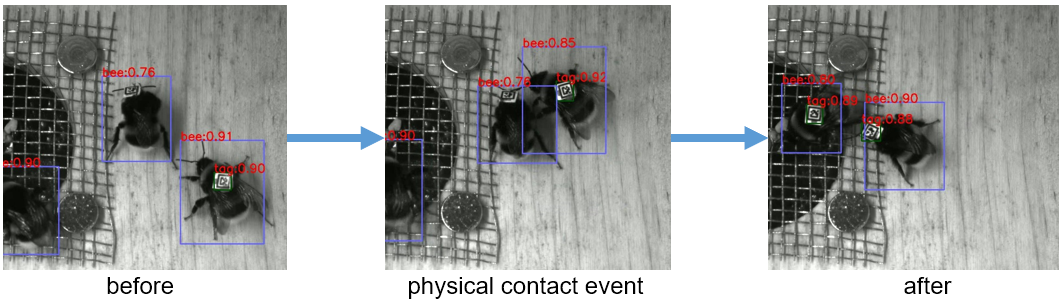

Extended Data Fig. 20 | During a physical contact event, the bounding boxes of two bumblebees are overlapped.
