## Supplementary Information for "Long-term exposure to microplastics and heat affects bumblebee behavior patterns, colony development and social networks"

† Contributed equally

### Supplementary Methods

#### Supplementary Method 1. Raman spectra analysis and PE identification

All spectra data were exported from WITec, and then processed and analyzed in MATLAB R2020b. We cropped the spectra from 500 to 3200 cm^-1^ which contained 1225 separate wavenumbers. We then used an automated algorithm (<https://github.com/michaelstchen/modPolyFit>) to remove the autofluorescence backgrounds^1^. Savitzky-Golay smoothing was then performed (polynomial order = 3, frame length = 11) on all spectra followed by normalization. Raman spectra of plastics feature peaks around particular wavenumbers, which varies across different plastic types ^2^. Principal component analysis (PCA) is commonly applied to reduce dimensionality of Raman spectrums and support microplastic identification^2,3^. We performed PCA on all spectra data and used the top 18 principal components (> 90% variance, Extended Data Fig. 2a) for K-means clustering.

The elbow method was used to determine the best K value. We varied K from 2 to 10 and iterated the clustering algorithm 1000 times for each K value. We calculated the sum of squared errors (SSE) of the target cluster containing all the known microplastics (Extended Data Fig. 2b). We set K value as 4 because this is the where the SSE curve starts to flatten out, meaning further increase of the number of clusters contributes little to clustering quality. K-means clustering was then iterated 3000 times on K = 4 to identify potential PE microplastic samples. The results were transformed into accumulation per mg (Extended Data Fig. 2c).

#### Supplementary Method 2. Relative effect sizes.

We used linear regression $model A$ to analyze time effects of microplastics and/or heat on a series of dependent variables. For each variable, there’s a natural trend over time, which is reflected in the coefficient estimate of the control group, $E_{c}\pm\sigma_{c}$. For each treatment group, we also have an affected trend over time, which is also reflected as the coefficient estimate, $E_{tr}\pm\sigma_{tr}$. We calculate the relative effect size of each treatment as

$$R_{tr}=\frac{E_{tr}-E_{c}}{\left| E_{c} \right|}\times100\%$$

Therefore, the sign of $R_{tr}$ reflects shows whether the treatment has a positive or negative impact, and the size of $\left| R_{tr} \right|$ reflects the degree of disturbance on the natural trend over time.

#### Supplementary Method 3. Estimation of microplastic accumulation in honeybees

Deng et al.^4^ collected honeybee samples from apiaries in China and measured in vivo microplastic particles. However, their work did not directly report the accumulation level. After sample collection, Deng et al.^4^ digested 0.5 g of honeybees and filtered through a 0.22 μm pore size membrane filter with 50 mm diameter. They randomly selected an area with 1 mm diameter and spotted more than 20 microplastics in the area. Therefore, we can estimate the accumulation level as

$$Acc>\frac{20 items\times\left( 50 mm \right)^{2}}{\left( 1 mm \right)^{2}\times0.5 g}=100\mathrm{items}/\mathrm{mg}.$$

#### Supplementary Method 4. Induced variation from two experimental iterations.

We ran the first experimental with 3 replicates (18 colonies) started on January 12th, 2024, and the second experiment with 2 replicates (12 colonies) started on June 5th, 2024. Since the two experimental iterations had different sample sizes and non-normal data distribution, we used Kruskal-Wallis tests to examine whether there’s induced variation from the two experimental iterations. We used iteration as categorization and performed Kruskal-Wallis tests for the dependent variables (see main text for details) that we used for effect analysis at colony, individual and social levels. This was realized by the kruskal.test() function in R. We also shuffled the 5 replicates into two random groups of 2 and 3 (9 possible combinations excluding the real categorization), and performed Kruskal-Wallis tests on these shuffled combinations. If the statistic values ($\chi^{2}$) of the real categorization are consistently higher than other shuffled combinations for variables across colony, individual and social levels, we confirm the existence of induced variation from two experimental iterations. The results are shown in Extended Data Fig. 14. For all dependent variables, the statistic value falls within the range of shuffled combinations, which means the two experimental iterations did not induce apparent variations to effect analysis at colony, individual and social levels.

#### Supplementary Method 5. Validation of nest structure stability

To improve the detection accuracy of colony nest structures of the first trial, we used the improved model on sample videos from the first trial, which were taken every 2.5 hours for each colony. Although the time span of sample videos (around 2×10^2^ frames per sample video, 2×10^3^ frames per day) is distinctly less than the time span of detection data, it still provides sufficient information to reconstruct the daily nest structures. We randomly selected a colony from the first trial (Extended Data Fig. 17a) and analyzed the variance of detected brood area and pot area against the accumulation of sampling frames. The values of detected brood area and pot area stabilized before the number of accumulated frames reached two thousand (Extended Data Fig. 17b). Therefore, the data from sample videos were sufficient to reconstruct the nest structures on a daily basis.
